# Fragile X Mental Retardation Protein regulates R-loop formation and prevents global chromosome fragility

**DOI:** 10.1101/601906

**Authors:** Arijita Chakraborty, Piroon Jenjaroenpun, Andrew McCulley, Jing Li, Sami El Hilali, Brian Haarer, Elizabeth A. Hoffman, Aimee Belak, Audrey Thorland, Heidi Hehnly, Chun-long Chen, Vladimir A. Kuznetsov, Wenyi Feng

**Affiliations:** Department of Biochemistry and Molecular Biology, SUNY Upstate Medical Univeristy, 750 East Adams Street, Syracuse, NY 13210, USA; Bioinformatics Institute, Agency for Science Technology and Research (A*STAR), 30 Biopolis Street, #07-01 Matrix, Singapore 138671; Institut Curie, PSL Research University, CNRS, UMR3244, F-75005 Paris, France; Sorbonne Université, F-75005 Paris, France; Department of Biology, Syracuse University, Syracuse, NY 13210, USA; Department of Urology, SUNY Upstate Medical University, 750 East Adams Street, Syracuse, NY 13210, USA; Department of Biomedical Informatics, University of Arkansas for Medical Sciences, Little Rock, Arkansas 72205, USA; Department of Biochemistry and Molecular Genetics, University of Virginia Health Systems, Charlottesville, VA 22908, USA; University of Maryland School of Medicine, 655 W. Baltimore Street, Baltimore MD 21201, USA

**Keywords:** Fragile X syndrome, *FMR1*, FMRP, *DHX9*, *UGT1A*, *CYP2C9*, R-loops, chromosome fragile sites, genome instability, xenobiotic metabolism

## Abstract

Fragile X syndrome (FXS) is the most prevalent inherited intellectual disability caused by mutations in the Fragile X Mental Retardation 1 (*FMR1*) gene. The protein product of *FMR1*, FMRP, is known as a translational repressor whose nuclear function is not understood. Here we report that FMRP is a genome maintenance protein. We show that FX cells exhibit elevated level of chromosome breaks, both spontaneous and replication stress-induced. We demonstrate that FMRP is required for abating R-loop accumulation, thereby preventing chromosome breakage. Through mapping FMRP-bound chromatin loci in normal cells and correlating with FX-specific chromosome breaks, we identified novel FXS-susceptible genes. We show that FX cells have reduced expression of the uridine diphosphoglucuronosyl transferase 1 family enzymes, suggesting defective xenobiotics glucuronidation and consequential neurotoxicity in FXS.

## INTRODUCTION

Fragile X syndrome (FXS) is responsible for the most common form of inherited intellectual disability and autism (Santoro et al., 2012). In most patients, FXS is caused by the (CGG)_n_ trinucleotide repeat expansion in the 5’-untranslated region of the Fragile X Mental Retardation (*FMR1*) gene located on Xq27.3, resulting in chromosome fragility at this locus, transcriptional silencing and loss of function of the gene product, FMRP (Coffee et al., 2002; Fu et al., 1991; Krawczun et al., 1985; Pieretti et al., 1991; Verkerk et al., 1991). FXS can also manifest as a result of mutations in the FMRP coding sequence, highlighting FMRP’s functional importance for the etiological basis for FXS (Ciaccio et al., 2017).

FMRP is an RNA binding protein and is estimated to bind ∼4% of the mRNAs in the brain and regulate their translation (Ashley et al., 1993). It is becoming increasingly clear that FMRP has multi-faceted functions. The best understood cellular function of FMRP is a translational repressor in the metabotropic glutamate receptor (mGluR)-mediated long-term depression (LTD) (Bear et al., 2004). The absence of FMRP permits increased level of protein synthesis at postsynaptic dendrites and prolonged LTD, which are normally inhibited by mGluR activation, thus causing many of the symptoms of FXS (Bear et al., 2004; Darnell et al., 2011; Nakamoto et al., 2007; Niere et al., 2012). Genome-wide studies have identified over 6,000 FMRP-interacting mRNAs, many of which are involved in synaptic signaling and function (Ashley et al., 1993; Brown et al., 2001; Darnell et al., 2011). However, only a small percentage of these putative FMRP targets have been validated by independent methods (Sethna et al., 2014) and mGluR antagonist drugs have yet to show efficacy in human patients despite preclinical success in animal models. A recent study also uncovered an enhancing role of FMRP in the translation of large autism-associated genes (Greenblatt and Spradling, 2018). Thus, it stands to reason that additional key FMRP targets that meet therapeutic potential remain at large. Consistent with its role as a translation repressor, FMRP is predominantly located in the cytoplasm and associates with the polysomes (Darnell et al., 2011; Khandjian et al., 2004). However, previous studies also demonstrated the involvment of FMRP in DNA damage response (Alpatov et al., 2014; Liu et al., 2012; Zhang et al., 2014).

In this study we set out to compare replication stress-induced chromosome breaks in lymphoblastoid cell lines derived from an individual with a full mutation at *FMR1* and an unaffected control and test the hypothesis that FMRP plays an important role in the maintenance of genome integrity. Lymphoblastoid cells have been used to reveal genetic basis for a range of neurological disorders including FXS (Brown et al., 2001). Using this system we demonstrate that the FX genome has increased susceptibility to chromosome breakage induced by replication stress, particularly at R-loop forming sites (RLFSs), where the RNA transcript hybridizes to homologous DNA on the chromosome, yielding a DNA:RNA hybrid and a displaced DNA single strand. Despite their many roles in normal cellular functions, R-loops can initiate conflicts between transcription and replication by creating a barrier to replication fork progression, causing chromosome breakage (Garcia-Rubio et al., 2018; Hamperl et al., 2017). We present evidence that FMRP is also a genome maintenance protein that prevents chromosome breakage by mediating R-loop accumulation, possibly through direct interaction with the gene substrates that are prone to R-loop formation during transcription and further enhanced by replication stress.

## RESULTS

### Fragile X cells show elevated DNA damage under replication stress

We chose lymphoblastoid cells derived from an individual diagnosed with FXS, who has a full mutation of *FMR1* (GM03200, henceforth “FX”), and an unaffected individual with a normal *FMR1* (GM06990, henceforth “NM”). The GM03200 cell line contains a reported 570 CGG repeat expansion at the 5’-UTR of *FMR1* (Hayward et al., 2016). We confirmed the repeat expansion by Southern blot analysis (∼590) and the lack of FMRP expression by western blot in the FX cell lines (Supplemental Fig. S1). We then analyzed genome instability in these cells by partially inhibiting replication with aphidicolin (APH, a DNA polymerase inhibitor) and causing a 10-20% increase of cells in S phase (Fig. 1A). Both cell lines showed dose-dependent increase of chromosome breaks upon APH treatment, evidenced by γH2A.X expression (a marker for DNA double strand breaks (DSBs)) using flow cytometry (Fig. 1B) and fluorescence microscopy (Supplemental Fig. S2). Compared to NM cells, FX cells demonstrated at least two-fold increase of APH-induced γH2A.X expression (Fig. 1B and Supplemental Fig. S2). Notably FX cells also showed more APH-induced γH2A.X foci per nucleus than NM cells did (Fig. 1C&D). These observations suggested that the absence of FMRP causes heightened susceptibility of FX cells to replication stress by APH. We next performed neutral Comet assay to investigate DSB formation at a single cell level. The results were consistent with the γH2A.X staining experiments where FX cells showed higher level of DSBs than their control counterparts (Fig. 1E). Additionally, the comet tail length distribution suggests that FX cells present comets with longer tails, often severed from the comet heads, indicating more severe DNA damage in the FX cells than in control cells (Fig. 1F). These results established that the FX genome is more prone to DNA damage upon replication stress than the control genome, thus prompting further investigation into the nature of genome instability in the FX cells.

**Figure 1.**
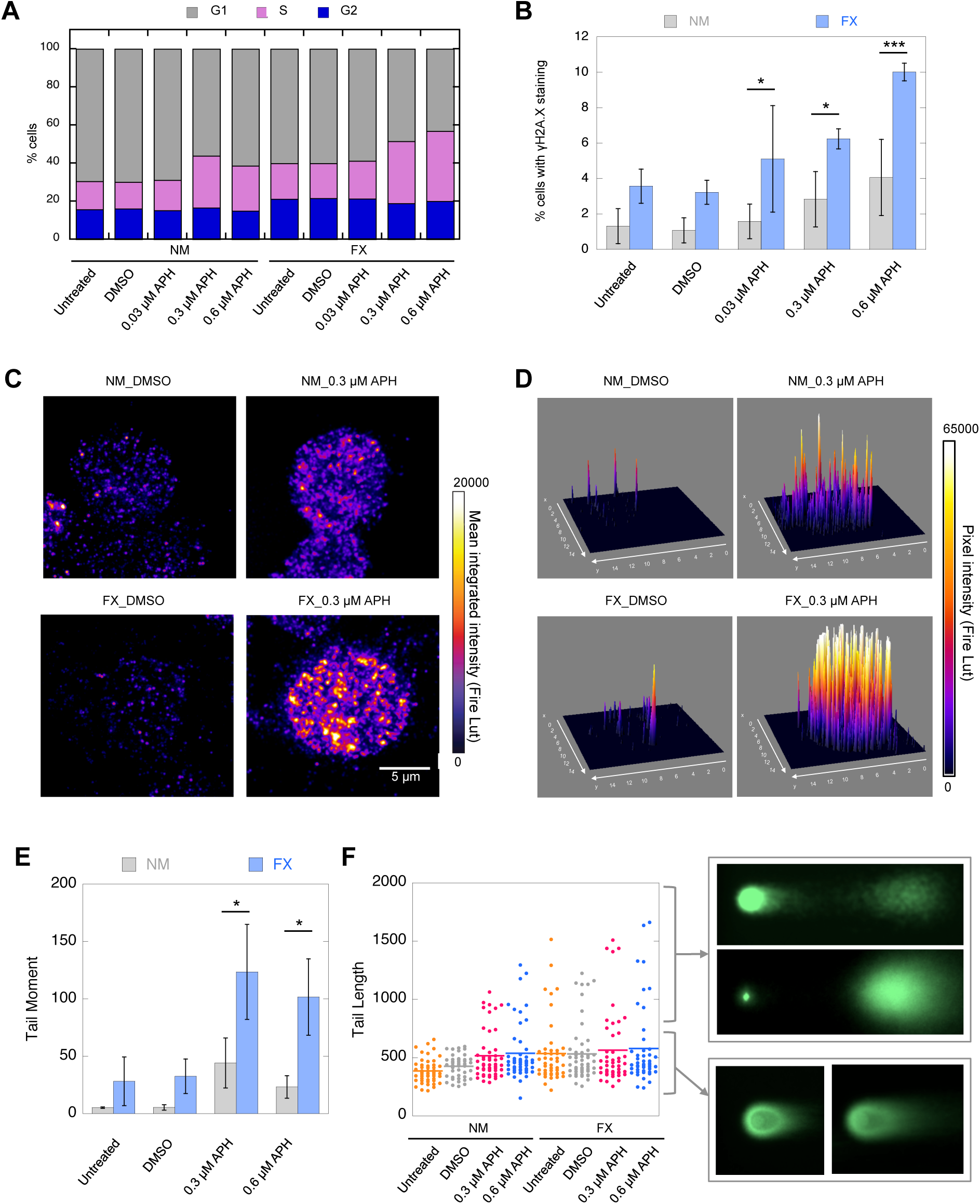
Fragile X cells show elevated DNA damage in vivo. (**A**) Cell cycle analysis by flow cytometry. Three independent experiments were performed and one representative experiment is shown. (**B**) Quantification of γH2A.X-positive cells by flow cytometic sorting. Ten thousand cells were analyzed in each of three independent experiments. Two-way ANOVA followed by Sidak’s multiple testing was performed. (**C**) Single cell view of γH2A.X foci distribution, expressed as mean integrated density (FIRE, LUT) for the indicated samples, in NM and FX cells. (**D**) 3D surface plots of γH2A.X intensity distribution in the cells from **(C**), with x and y axes in microns and the z axis in pixel intensity after thresholding at each focus (FIRE, LUT), in NM and FX cells. (**E&F**) Single-cell analysis of DNA damage by Comet assay. Three biological replicates were performed (n>50 in each replicate) and all trended similarly. A representative experiment is shown for Tail Moment measurement. One-way ANOVA followed by Sidak’s multiple testing was performed. Error bars denote standard errors of mean. Tail Lengths from one representative experiment are shown in box plot (F). Representative images of comets with short or long Comet tail length are shown.

### Genome-wide DSB mapping by Break-seq

Here we adapted Break-seq, a powerful technology we first developed in yeast (Hoffman et al., 2015), to the mammalian system and mapped genome-wide chromosome breaks in untreated cells and cells treated with APH or with equal volume of the vehicle, dimenthyl sulfoxide (DMSO) (Fig. 2A). Break-seq data quality check and control experiments are described in Supplemental Information and shown in Supplemental Table S1 and Supplemental Fig. S3. For each strain/treatment combination, *e.g.*, “FX_0.03 µM APH”, *consensus DSBs* from at least two replicate experiments, regardless of the total number of replicates, were derived (Supplemental Table S2). The DSBs from 0.03 µM and 0.3 µM APH-treated samples were further pooled into a composite dataset of “APH-treated DSBs”, for each cell line, followed by comparison with the DMSO-treated control to identify DSBs shared by DMSO- and APH-treatment as well as those specific to each treatment (Supplemental Fig. S4A&B, Supplemental Table S2).

**Figure 2.**
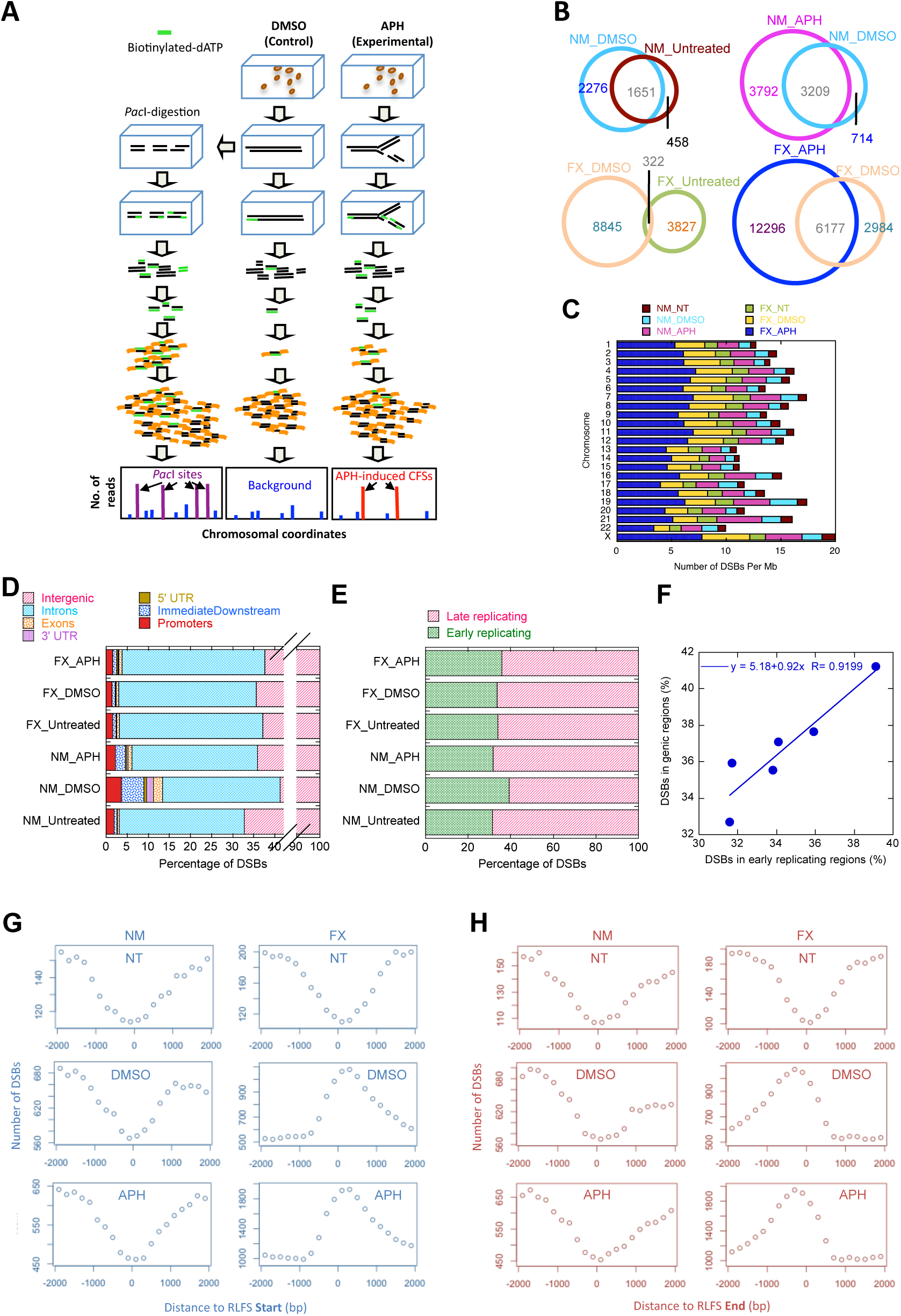
Break-seq mapping shows high level of DSBs in FX cells and drug-induced DSBs in FX correlate with RLFSs. **(A)** Break-seq methodology. (**B**) Chromosomal distribution of the number of DSBs per Mb of DNA in the indicated categories. **(C)** Venn diagrams showing the concordance between samples of different treatments. FX cells show “reprogramming” of drug-induced DSBs compared to those in the untreated cells. **(D)** Distribution of DSB peaks relative to genes in the indicated samples. Genic features include introns, exons, 5’- and 3’-UTRs, promoters, and the immediately downstream (<1 kb from the 3’-UTR) regions (ImmediateDownstream). (**E**) Distribution of DSBs in early vs. late replicating regions of the genome, as defined by Hansen RS et al, in the indicated samples. See Methods for details. (**F**) Correlation between the percentage of DSBs associated with genes and the percentage of DSBs with early replication timing. Aggregated DSBs from the indicated samples around the start **(G)** or the end **(H)** of R loop forming sites (RLFSs) in a 4000 bp window centering on the RLFS.

### FX cells show global increase of DSB formation and abnormal response to drug treatment

In all experiments, FX cells produced 2-2.5 fold more DSBs than NM cells with or without drugs (Fig. 2B), consistent with high levels of DNA damage observed in FX cells. The X chromosome showed the highest density of DSBs in FX cells (Fig. 2C). Overall, there was a higher concordance of DSBs in NM cells than FX cells: while 78% of the DSBs in untreated NM cells were also present in the DMSO-treated NM cells, there was only an 8% concordance between untreated and DMSO-treated FX cells (Fig. 2B). This was unlikely an artifact because 1) the vast majority of DSBs (83%) in untreated NM cells were also identified in untreated FX cells (Supplemental Fig. S4C); and 2) 67% of the DSBs found in DMSO-treated FX cells were also identified in APH-treated FX cells (Fig. 2B). Overall, 83%, 16% and 62% of DSBs in the untreated, DMSO-treated, and dual-treated NM cells, respectively, were found in the corresponding treatments of FX cells (Supplemental Fig. S4C). We concluded the following. First, spontaneous DSBs were largely concordant between NM and FX cells. Second, spontaneous DSBs in NM cells remained in DMSO treatment, whereas FX cells “reprogrammed” DSBs in response to DMSO, suggesting unique chemical-genetic interactions in drug-treated FX cells, as subsequent analysis would reveal. Third, the dual treatment with DMSO and APH both enhanced existent DSBs (from DMSO treatment alone) and induced new DSBs, in both cell types.

### DSBs are correlated with late replication timing or drug-induced transcription

Approximately 30 to 40% of DSBs in all samples occurred in genic regions with a dominant presence in the introns. In NM cells genic association of DSBs increased from 33% with spontaneous breakage to 41% with DMSO treatment, followed by a return to 36% with dual drug treatment (Fig. 2D). In contrast, FX cells showed higher level (37%) of DSB-gene association than NM cells in the untreated condition, but remained constant with 36% and 38% in DMSO-treated and dual-treated conditions, respectively (Fig. 2D). These results suggested FX cells might be more transcriptionally active than NM cells, whereas NM cells might have stronger transcriptional response to drug treatment than do FX cells. Supporting this notion it was shown that FMRP knock-out mouse neurons exhibit increased gene expression compared to control neurons (Korb et al., 2017). Thus, we surmised that the gene-associated DSBs were transcription-related and sought mechanisms for intergenic DSB formation. Delayed replication timing is one of the hallmarks for APH-induced chromosome fragility (Hellman et al., 2000; Le Beau et al., 1998; Palakodeti et al., 2004; Wang et al., 1999). Using published Repli-seq data of the NM cells (Hansen et al., 2010) we divided the genome into 50-kb early- or late-replicating segments and calculated the percentage of DSBs in each segment (Supplemental Information, Supplemental Methods and Discussion). All six samples showed enrichment of DSBs in late-replicating regions (p < 10E-3) (Fig. 2E). Notably, the percentage of DSBs that fall in the early replicating regions is correlated with the percentage of DSBs within genes (Fig. 2F). Together these results suggested two main types of chromosome breakage—those in the intergenic regions that undergo delayed replication and those in the gene-rich early replicating regions that experience elevated level of gene transcription.

### Preferential association between drug-induced DSBs and R-loop forming sequences in FX cells

If the gene-associated DSBs were due to drug-induced replication-transcription conflict at actively transcribing genes, DSBs might correlate with R-loops, which are co-transcriptional structures. We surveyed a database of R-loop forming sequences (RLFSs) predicted by a previously described algorithm (Wongsurawat et al., 2012) for correlation with DSBs (Supplemental Table S3). Untreated cells did not show enrichment of spontaneous DSBs at RLFSs; however, DMSO-treated NM and FX cells both showed significant enrichment of DSBs at RLFSs. Notably, DSBs in APH-treated NM cells were no longer enriched at RLFSs whereas those in APH-treated FX cells remained associated with RLFSs. These results were corroborated by the absolute distance measurements between DSBs and RLFSs (Table 1) using GenometriCorr (Favorov et al., 2012). We concluded that 1) DMSO elicits transcriptional response, possibly through oxidative stress (Supplementary Information), in both NM and FX cells and cause DSBs at RLFSs within actively transcribing genes; 2) replication inhibition by APH triggers NM cells to deploy a mechanism to protect genes from DSBs at RLFSs, whereas FX cells lacked such a mechanism. Consistent with our interpretation, APH-specific DSBs in FX cells showed the greatest association with genes compared to DSBs in any other category (Supplemental Table S4). Aggregated distribution of DSBs around RLFSs showed an enrichment of DSBs immediately downstream of the RLFS start as well as immediately upstream of the RLFS end, specifically in the drug-treated FX cells (Fig. 2G&H). Overall, these results led us to conclude that FX cells form DSBs at RLFSs when treated with DMSO, a response that is further enhanced by APH. The correlation between DSBs and RLFSs is the strongest on chromosomes 1, 13, 14, 15, 21, and 22, which contain ribosomal DNA (rDNA) clusters, followed by chromosomes 2, 6, and 12 (Table 1). Notably, the DSBs on the rDNA-bearing chromosomes were not confined to the rDNA loci. We also compared the DSBs to a composite list of DRIP-seq (DNA:RNA hybrid Immunoprecipitation followed by sequencing) signals, generated by merging all DRIP-seq signals in NT2 and K562 cell lines to minimize cell type-specific differences (Sanz et al., 2016). The results largely recapitulated the comparison between DSBs and RLFSs (Supplemental Table S3). Because RLFS prediction is an unbiased approach, we focused on testing if RLFSs are more susceptible to breakage in FX than in NM cells.

**Table 1.** Statistical analysis of absolute distance between RLFSs and DSBs in the indicated categories on each chromosome using GenometriCorr. Shown are the projection test p-values (P) and projection test observed to expected ratios (R). Samples with R values greater than 3.5 are shown in red.

| Chr | FX.Untreated |  | FX.DMSO |  | FX.APH |  | NM.Untreated |  | NM.DMSO |  | NM.APH |  |
| --- | --- | --- | --- | --- | --- | --- | --- | --- | --- | --- | --- | --- |
|  | P | R | P | R | P | R | P | R | P | R | P | R |
| 1 | 0.355 | 1.041 | 2.22e-16 | 3.799 | 1.07e-14 | 2.764 | 0.224 | 1.239 | 0.086 | 1.514 | 0.353 | 0.816 |
| 2 | 0.314 | 0.644 | 2.21e-11 | 3.625 | 1.55e-10 | 2.629 | 0.236 | 1.212 | 1.73e-06 | 3.797 | 0.010 | 1.850 |
| 3 | 0.446 | 0.691 | 1.29e-06 | 3.028 | 1.81e-07 | 2.579 | 0.329 | 0.000 | 0.024 | 2.283 | 0.131 | 1.430 |
| 4 | 0.283 | 0.398 | 2.39e-05 | 3.008 | 1.44e-10 | 3.227 | 0.355 | 0.801 | 0.036 | 2.209 | 0.199 | 1.281 |
| 5 | 0.217 | 0.348 | 1.09e-06 | 3.244 | 1.63e-08 | 2.799 | 0.244 | 0.000 | 0.007 | 2.553 | 0.148 | 1.382 |
| 6 | 0.090 | 0.000 | 6.36e-07 | 3.746 | 5.08e-04 | 2.065 | 0.336 | 0.000 | 0.134 | 1.533 | 0.028 | 0.000 |
| 7 | 0.489 | 0.868 | 6.30e-06 | 2.515 | 4.01e-10 | 2.575 | 0.108 | 0.000 | 0.104 | 1.483 | 0.126 | 1.339 |
| 8 | 0.150 | 0.298 | 4.34e-06 | 2.965 | 1.06e-04 | 2.085 | 0.182 | 1.362 | 0.203 | 1.294 | 0.131 | 0.407 |
| 9 | 0.301 | 0.413 | 3.91e-07 | 3.217 | 4.19e-12 | 3.197 | 0.174 | 1.392 | 0.016 | 2.174 | 0.109 | 0.461 |
| 10 | 0.137 | 0.289 | 2.85e-07 | 3.279 | 3.85e-09 | 2.806 | 0.302 | 1.041 | 0.088 | 1.638 | 0.373 | 1.012 |
| 11 | 0.070 | 0.233 | 7.45e-05 | 2.213 | 0.001 | 1.643 | 0.118 | 0.000 | 0.047 | 1.708 | 0.123 | 0.528 |
| 12 | 0.044 | 0.000 | 4.91e-11 | 4.040 | 7.32e-09 | 2.743 | 0.209 | 0.000 | 0.076 | 1.703 | 0.350 | 1.040 |
| 13 | 0.372 | 0.000 | 9.55e-08 | 6.960 | 8.71e-10 | 5.431 | 0.347 | 0.000 | 0.482 | 0.000 | 0.260 | 0.000 |
| 14 | 0.190 | 1.333 | 8.41e-06 | 3.516 | 6.05e-11 | 3.813 | 0.035 | 2.782 | 5.87e-04 | 4.219 | 0.103 | 1.620 |
| 15 | 0.109 | 1.655 | 3.62e-07 | 4.231 | 2.64e-11 | 3.926 | 0.377 | 0.000 | 0.054 | 1.962 | 0.088 | 1.636 |
| 16 | 0.050 | 0.214 | 0.00e-00 | 5.667 | 0.00e-00 | 4.522 | 0.484 | 0.769 | 0.060 | 1.626 | 0.291 | 1.112 |
| 17 | 0.056 | 0.221 | 2.62e-08 | 3.289 | 7.21e-13 | 3.158 | 0.322 | 0.434 | 0.353 | 1.048 | 0.165 | 1.273 |
| 18 | 0.103 | 1.786 | 1.14e-04 | 3.962 | 3.54e-04 | 2.763 | 0.144 | 1.498 | 0.061 | 2.238 | 0.237 | 1.191 |
| 19 | 0.057 | 0.459 | 2.32e-04 | 1.941 | 2.88e-07 | 1.916 | 0.162 | 0.446 | 0.429 | 0.999 | 0.065 | 0.635 |
| 20 | 0.456 | 0.803 | 5.62e-04 | 2.831 | 9.08e-08 | 3.164 | 0.135 | 1.568 | 0.039 | 2.140 | 0.218 | 1.239 |
| 21 | 0.230 | 0.000 | 5.01e-09 | 7.108 | 2.22e-13 | 6.001 | 0.426 | 0.000 | 0.014 | 2.812 | 0.432 | 0.895 |
| 22 | 0.093 | 0.000 | 1.80e-09 | 5.343 | 3.99e-10 | 3.764 | 0.462 | 0.646 | 0.016 | 2.142 | 0.009 | 2.158 |
| X | 0.184 | 1.351 | 0.081 | 1.482 | 0.098 | 1.339 | 0.113 | 1.636 | 0.024 | 2.132 | 0.077 | 1.555 |
| ALL | 5.47e-06 | 0.528 | 0 | 3.162 | 0 | 2.684 | 0.041 | 0.715 | 8.93e-14 | 1.976 | 0.150 | 1.088 |

### Ectopic expression of FMRP, but not the FMRP-I304N mutant, reduces RLFS-induced DSBs in the model organism *Saccharomyces cerevisiae*

We employed a modified yeast-based recombination assay (Prado and Aguilera, 2005) to measure DSB frequency resulting from programmed transcription-replication conflict induced by human RLFSs (Fig. 3A). In the absence of RLFS insertion there was a 4.7-fold enhancement of recombination frequency (RF) on the plasmid with convergent replication and transcription compared to a codirectional configuration (Supplemental Fig. S5A), consistent with the previous observation (Prado and Aguilera, 2005). Two human RLFSs, when inserted in the sense direction, each caused elevated RF over the control sequence (non-RLFS), in the convergent replication-transcription configuration specifically (2 and 4 fold, p=0.0024 and p<0.0001, respectively, Supplemental Fig. S5B). RF was further enhanced in a strain lacking RNase H1, an enzyme known to resolve R-loops by degrading the RNA:DNA hybrid: ∼2 and 1.5-fold for sense and anti-sense orientation, respectively, for RLFS-1; and ∼1.2 fold for both sense and anti-sense orientations, for RLFS-2 (Supplemental Fig. S5B&C). Because RLFS-2 already induced high RF, further enhancement by eliminating RNaseH1 was only moderate. Next we asked if ectopic expression of FMRP would decrease RLFS-induced DSBs. Expression of empty vector did not alter the RLFS-induced RF (comparing Supplemental Fig. S5B to Supplemental Fig. S5D). A significant drop in RF was observed for expression of FMRP (2 and 1.6 fold for RLFS-1 and RLFS-2, respectively) or a positive control, RNaseH1, but not for a non-specific RNA binding protein She2, compared to empty vector (Fig. 3B). Finally, a mutant FMRP containing an I304N substituion in the KH2 domain, a rare *de novo* mutation that led to FXS (De Boulle et al., 1993; Siomi et al., 1994), no longer suppressed RF (Fig. 3B). We also verified that FMRP and FMRP-I304N showed similar level of protein expression in yeast (Supplemental Fig. S5E). The I304N mutation abolishes FMRP binding to mRNA and the polysome (Feng et al., 1997). Our results thus suggest that the KH2 domain is also involved in interaction with R-loops.

**Figure 3.**
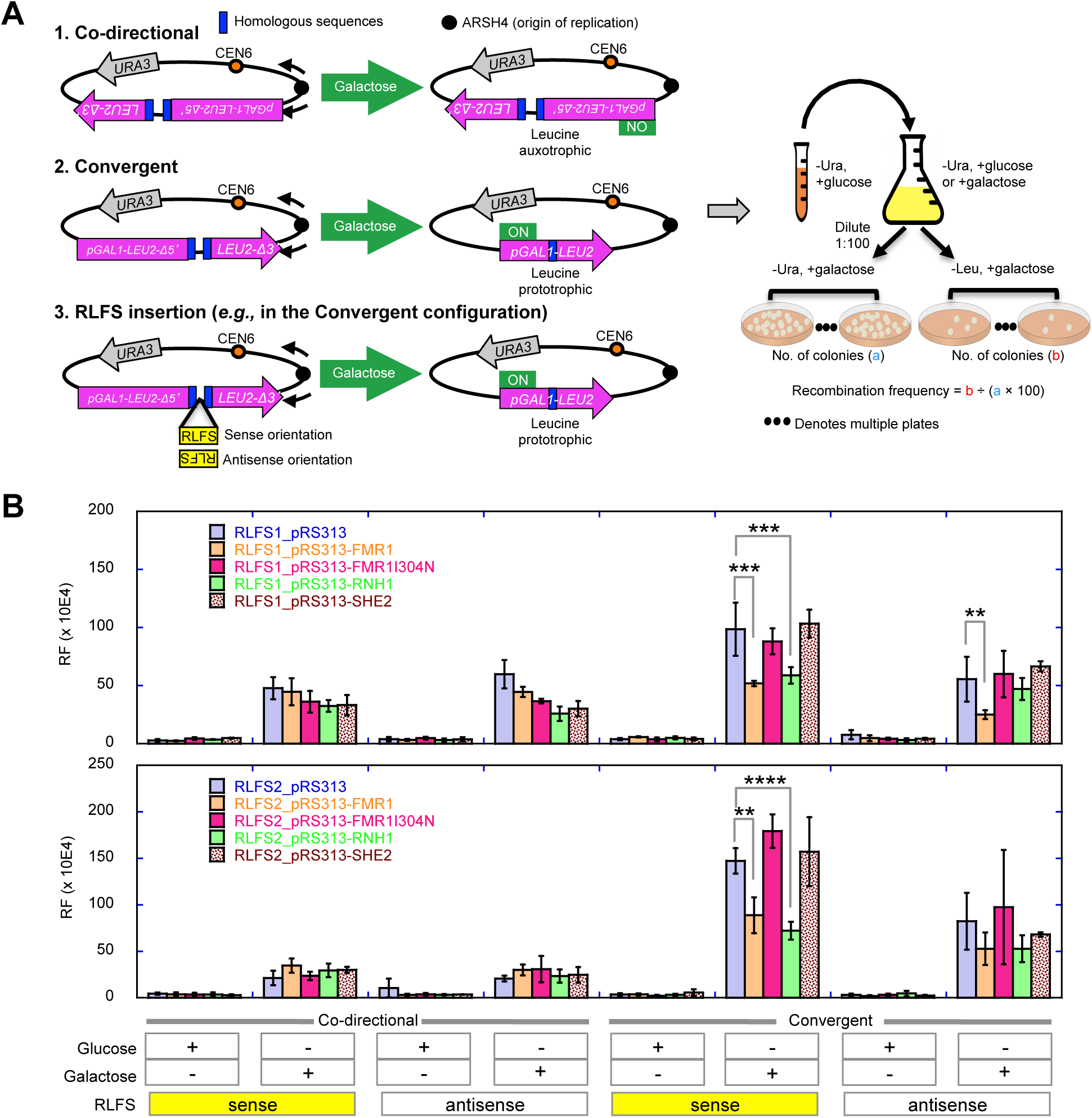
FMRP expression suppresses RLFS-induced DSB formation. (**A**) A non-functional *LEU2* marker containing two inserted direct repeats and driven by a galactose-inducible *GAL1* promoter was placed next to an origin of replication (*ARSH4*) such that the direction of transcription is convergent or codirectional with respect to the direction of the proximal replication fork. Upon galactose induction, convergent replication and transcription would induce DSBs and homologous recombination repair to generate a functional *LEU2*, resulting in leucine prototrophy. Two RLFSs from the human genome (RLFS1-1 from the promoter of *FMR1* and RLFS-2 from intron 5 of Fragile Histidine Triad) were inserted between the direct repeats to test for enhanced DSB and recombination. A non-RLFS sequence without predicted R-loop forming propensity and with similar G-richness in both strands served as control. All sequences were similar in size (∼500 bp). The RLFSs were inserted in the sense or anti-sense orientation with respect to *LEU2* transcription (*i.e.*, G-rich strand on the non-template or template strand, respectively), with the sense orientation expected to preferentially induce R-loop formation. The control sequence was also inserted in two orientations and no difference in RF was observed between them (Fig. S5C). RF is calculated based on the percentage of leucine prototrophs after plating. (**B**) The effect of ectopic expression of indicated genes on the pRS313 plasmid, under the CMV promoter, on RLFS-induced RF. See Figure S5 for additional control experiments.

### FMRP co-localizes with RNA:DNA hybrid during replication stress

Our study so far suggests that FMRP binds to its mRNA targets co-transcriptionally and prevents stable R-loop formation. A recent study reported that FMRP localizes to the nucleus upon APH treatment in mouse embryonic fibroblasts (Alpatov et al., 2014). We asked if the human FMRP similarly alters its subcellular distribution in response to APH. The total level of FMRP remained constant throughout the treatments (Fig. 4A&B). In contrast, the nuclear fraction of FMRP increases with APH treatment compared to the vehicle control, with the highest level of nuclear FMRP upon 0.3 μM APH treatment (Fig. 4A&C). To further demonstrate that FMRP is associated with R-loops we examined their co-localization by immunostaining. In untreated cells, co-localization of FMRP and RNA:DNA hybrid was relatively confined to the nuclear periphery and cytoplasm. With drug treatment, the co-localization became enriched in the nucleus (Fig. 4D). Three dimensional volumetric reconstruction further confirmed the co-localization (Supplemental Movie S1). We then asked if FMRP interacts with proteins known to participate in RNA:DNA hybrid resolution. To date we have detected interaction between FMRP and DHX9 (Fig. 4E), a helicase that has been shown to suppress R-loop formation and prevents chromosome breakage (Cristini et al., 2018).

**Figure 4.**
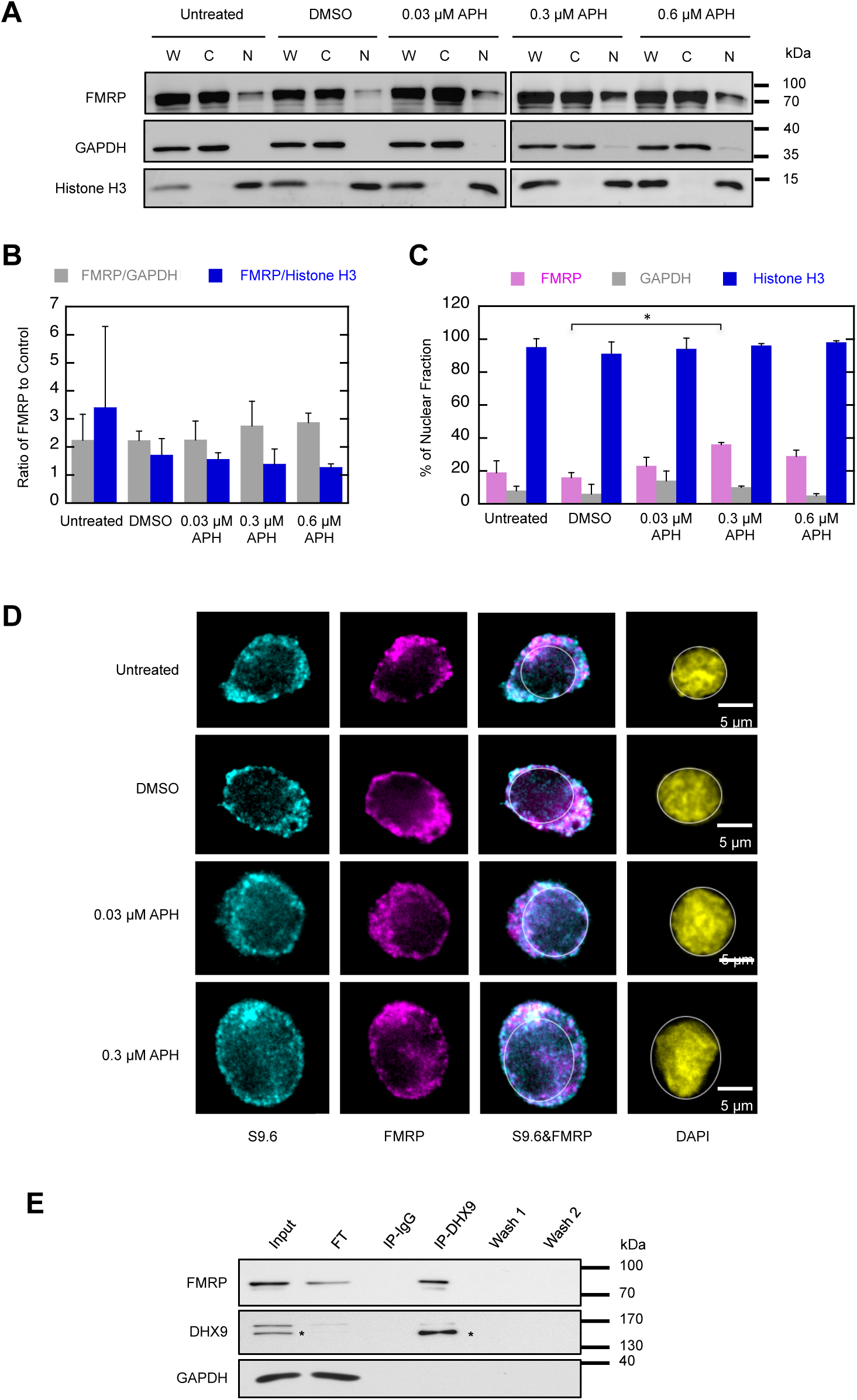
FMRP is enriched in the nucleus upon replication stress, co-localizes with RNA:DNA hybrids and co-immunoprecipitates with DHX9. (**A**) Western blots of FMRP in the nuclear (“N”) and cytoplasmic (“C”) fractions. “W” (whole cell extract) from the same number of cells used for fractionation is also shown for reference. Two independent experiments were performed and a representative experiment is shown. GAPDH and Histone H3 serve as cytoplasmic and nuclear controls, respectively. (**B**) Total FMRP level expressed as ratio of FMRP over control proteins (GAPDH or Histone H3) in the whole cell extracts (n=3). (**C**) Percentage of nuclear fraction of FMRP expressed as the percentage of the band intensity for “N” over that of the sum of “N” and “C” for each sample (p = 0.04, one-way ANOVA followed by Tukey’s multiple tests). (**D**) Co-localization of FMRP and RNA:DNA hybrids. Immunofluorescence images of untreated, DMSO- and APH-treated NM cells co-stained for RNA:DNA hybrids (cyan), FMRP (magenta) and nucleus (yellow, outlined). Immuno-staining is shown in a single Z-plane. Scale bar, 5 µm. (**E**) Co-immunoprecipitation of FMRP and DHX9 using the anti-DHX9 to pull down DHX9 and immunoblotted for both FMRP and DHX9. GAPDH served as negative control. The asterisks indicate the lower band of a doublet signal in is the DHX9 protein, which is present in the DHX9-immunoprecipitated complex and absent in the IgG-precipitated control complex (“IP-IgG”).

### FMRP chromatin binding sites are associated with RLFSs

We predicted that FMRP interacts with its mRNA substrates on the chromatin during transcription prior to transporting them into the cytoplasm for translation. We tested this hypothesis by performing a ChIP-seq experiment to identify FMRP chromatin-binding sites. First, we demonstrated the specificity of the anti-FMRP antibody in immunoprecipitation (Supplemental Fig. S6A). Using this antibody we compared the ChIP-seq signals from the control cell line to those from the FX cell line and identified 5238 sites that were enriched in the control cell line as putative FMRP chromatin-binding sites (Fig. 5A). Among these FMRP-binding sites, 54.8% are located in the genic regions, predominantly in the introns (44.7%) (Supplemental Fig. S6B). We then used ChIP-qPCR to validate select top FMRP-binding genes (*DLG1, CLNK, ASTN2* and *ANK1*) by way of certifying the efficacy of ChIP-seq methodology (Supplemental Fig. S6C). We also validated those genes that have been previously shown as mRNA substrates for FMRP and contain both FMRP-binding sites and APH-induced DSBs in FX cells: *GRIA1, GRM5, MTOR, PTEN* and *UGT1A* (Supplemental Fig. S6C). At a first pass, surprisingly only 283 FMRP-binding sites in 191 genes overlapped with an RLFS (Fig. 5B). FMRP-binding sites with overlapping RLFSs also showed relatively lower ChIP signals than stand-alone FMRP-binding sites (Supplemental Fig. S6D), indicating genes where these two features overlap are under-represented in the genome (p < 2.2e-16, Fisher’s exact test). However, the absolute distance between FMRP-binding sites and RLFSs was significantly shorter than expected (Fig. 5C). Moreover, FMRP-binding sites with overlapping RLFSs are enriched at the promoter and transcription start site (Fig. 5B), a pattern that was not observed for stand-alone FMRP-binding sites. Therefore, we concluded that FMRP-binding sites tend to be adjacent to, rather than overlapping with, RLFSs. We then honed in on 487 genes harboring overlapping or adjacent (< 1 kb apart) FMRP bindings sites and RLFSs (Supplemental Table S5). Expression of these genes was enriched in the brain (p = 5.7E-2) and particularly, in amygdala (p = 3.0E-2). There was also a moderate enirchment of genes in “learning or memory” (p = 6.7E-1, Supplemental Fig. S6E). Finally, 16 of 36 previously validated mRNA substrates for FMRP (Sethna et al., 2014) were identified as its chromatin-binding sites and/or APH-induced DSBs in FX cells, with 4 genes (*GRIA1, GRM5, MTOR*, and *PTEN*) containing both (Supplemental Fig. S6F), consistent with the roles of glutamate receptors and mTOR signaling in FXS.

**Figure 5.**
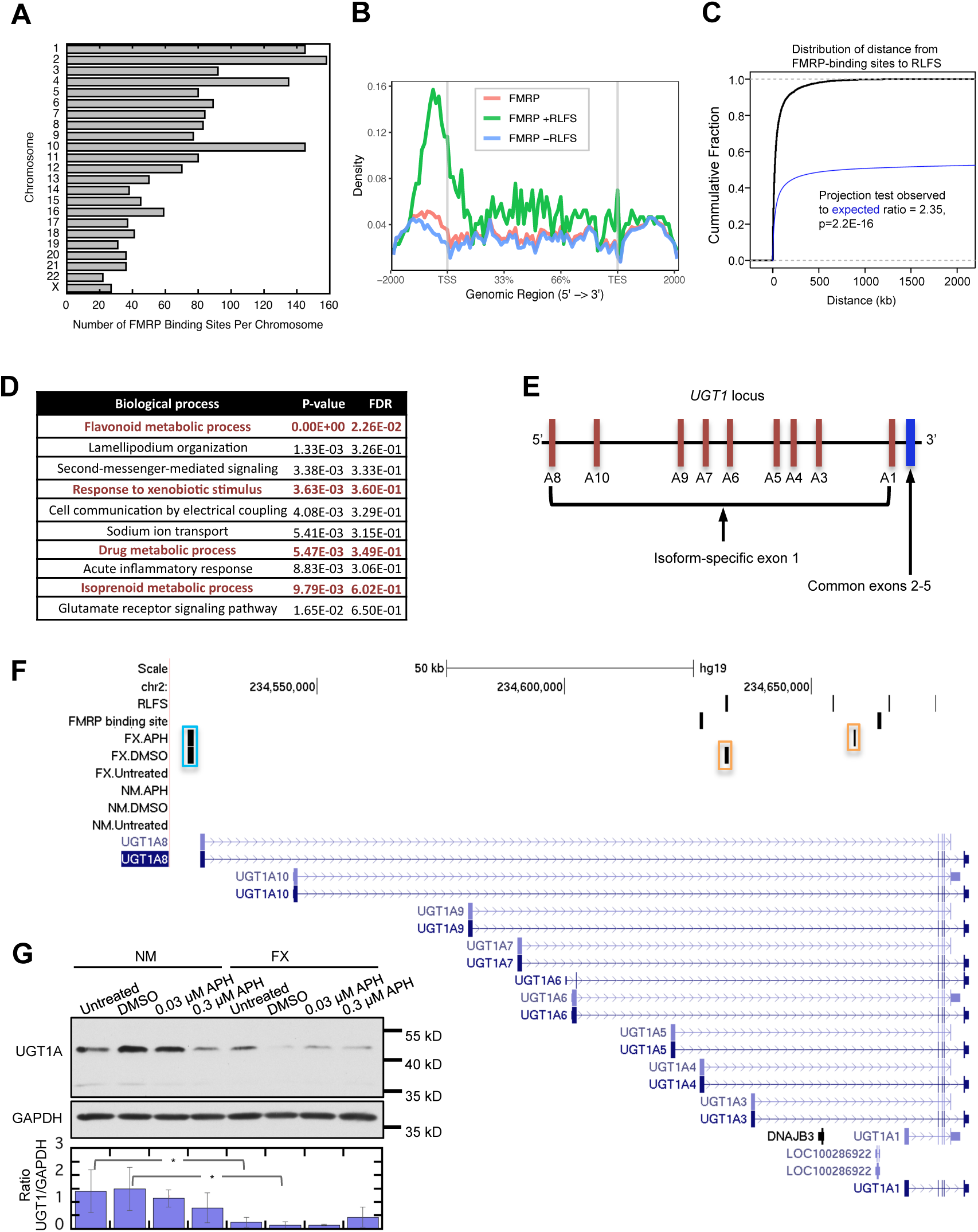
Analysis of chromatin binding sites of FMRP. (**A**) Distribution of the number of FMRP binding sites (FBSs) per chromosome. (**B**) Distribution of the FBSs inside genic regions, and in regions upstream of TSS (Transcription Start Sites) and TES (Transcription End Sites). Red, green and blue lines represent distribution of 5238 FBSs, 283 FBSs with overlapping RLFS and 4955 FBSs without overlapping RLFS, respectively. (**C**) Observed absolute distance between FBSs and RLFSs (blue) compared to the expected distance if uncorrelated (black line). (**D**) Biological processes derived from WebGestalt (WEB-based GEne SeT AnaLysis Toolkit) that are enriched for those genes bound by FMRP; FDR, false discovery rates. (**E**) Schematic representation of UGT1A subfamily alternative splicing isoforms. (**F**) UCSC genome browser screen shot of FX-specific DSBs shown as vertical bars (upstream of the gene, cyan box; in the intron preceding the common exons 2-5, orange boxes) in *UGT1* family genes. (**G**) Representative Western blots of UGT1 protein expression with GAPDH as control and quantification using ratio of UGT1 to GAPDH derived from three biological replicates. Error bars denote standard deviation. Two-way ANOVA followed by Sidak’s multiple testing were performed.

### Identification of novel FXS-associated genes with coalesced sites of DSBs and FMRP-binding sites

We reasoned that the FMRP-binding genes may be at heightened risk for DSBs in FX cells and identifying these genes can lead to discovery of novel FXS-associated genes. First, we annotated the genes neighboring spontaneous DSBs in NM cells (5 kb maximum distance) and found no significant (p < 0.001) GO (Gene Ontology) enrichment. In contrast, spontaneous DSB-associated genes in FX cells were enriched in “neuron projection development”, “synapse organization” and “neuron cell-cell adhesion” (Supplemental Table S6). Genes containing overlapping RLFSs and drug-induced DSBs in the FX cells were enriched in polysaccharide metabolism, including flavonoid glucuronidation and membrane organization pathways (Supplemental Table S6). Interestingly, 3993 genes associated with the 5238 FMRP-binding sites were also enriched in flavonoid/xenobiotics metabolism (p = 0; Fig. 5D). The end step of phase II xenobiotics metabolism is glucuronidation catalyzed by the uridine diphosphoglucuronosyl transferases (UGTs) comprised of the UGT1 and UGT2 families. The UGT1 family is derived from a single gene locus through alternative splicing and joining of an isoform-specific exon 1 with four common exons 2-5 (Fig. 5E). The *UGT1* family, and not *UGT2*, contain co-localized FMRP-binding sites and FX cells-specific DSBs, the latter of which can potentially impact the expression of all isoforms by virtue of residing in the intron preceding the common exons 2-5 (Fig. 5F). Only *UGT1A1* is possibly spared because the DSBs precede exon 1 sequence for *UGT1A1.* Two phase I xenobiotics metabolic genes, *CYP2C9* and *CYP2C19*, also contain FX-specific and drug-induced DSBs (Supplemental Fig. S7A&B). We report that untreated FX cells showed ∼2 fold reduction of UGT1 expression compared to NM cells, and greater reduction was seen in drug-treated FX cells (Fig. 5G). Similarly, FX cells showed reduction of CYP2C9 expression (Supplemental Fig. S7C).

### The FX lymphoblast-associated phenotypes are recapitulated in FX fibroblasts

So far all our experiment had been done with lymphoblast cell lines. To ensure that the observed FX-associated phenotypes were not due to a tissue-specific artifact, we performed select experiments with fibroblasts derived from a FX individual, GM05848, which contains a full mutation of 700 CGG repeats at *FMR1* (Sheridan et al., 2011) and from a sex- and age-matched unaffected control (GM00357). Souther blot analysis revealed ∼730 repeat expansion in the FX individual (Supplemental Fig. S1). We confirmed that FX fibroblasts also showed increased DNA damage by γH2A.X staining, particularly during APH treatment (Supplemental Fig. S8A&B). This was accompanied by the observation that FX fibroblasts also showed increased RNA:DNA foci formation in the nucleus upon drug treatment (Supplemental Fig. S8C), consistent with increased R-loop formation in these cells. Moreover, the fibroblast cells—with their larger nuclei— enabled the detection of RNA:DNA foci localizing to what appeared to be dark areas of the nucleus, possibly the nucleoli (Supplemental Fig. S8C). Finally, we also observed decreased UGT1 expression in the FX fibroblasts by western blots (Supplemental Fig. S8D).

## DISCUSSION

### FMRP as an R-loop regulator and a guardian of genome integrity

The main discovery from our study is the genome-wide chromosomal breakage in FX cells, with or without replication stress, suggesting FMRP is a guardian of the genome. This represents a novel development in our understanding of the FXS etiology. FX cells were defined by the detection of a fragile site named FRAXA at the *FMR1* locus, specifically induced by folate stress, in individuals with full mutation of the CGG repeat expansion. While there were an abunance of studies measuring FRAXA site expression, relatively few compared the number of common fragile sites (CFSs), which are induced by APH, in FX cells to controls. One study reported more than 3 fold increase of the rates of CFSs in FX patients (27.9%) compared to unaffected controls (7.9%) when treated with APH (Murano et al., 1989). However, the authors did not emphasize this difference and concluded that age difference between the test and control groups may have confounded the results. We surmise that this was an unexpected finding which led to the downplayed conclusion. We also note that these earlier studies were based on a cytological screening method with low resolution and sensitivity, guided by primary focus on detecting FRAXA in the FX cells. Thus, it is owing to the Break-seq technology with its unparalleled sensitivity that the detection of global DSB formation in the FX genome was enabled, a testament to the utility of Break-seq in other disorders with underlying etiological basis of genome instability. The global induction of DSBs in the FX genome was corroborated by increased γH2A.X stainining. This observation apparently differs from a previous study reporting decreased γH2A.X staining in *fmr1^-/-^* mouse embryonic fibroblasts (Alpatov et al., 2014). As the mutation in the FX individual is repeat expansion-induced epigenetic silencing of *FMR1*, whereas the mutation in the mouse model involves the deletion of exon 5 of *FMR1*, we suggest the apparent discrepancy in our observations at least partially stemmed from these key differences in our experimental systems. Indeed, this apparent discrepancy is echoed by opposing observations in the dFMRP1-deficient *Drosophila* model of FXS (Liu et al., 2012; Zhang et al., 2014), wherein Zhang et al. reported decreased γH2A.X foci formation when cells were treated with hydroxyurea and Liu et al. demonstrated hypersensitivity to genotoxic chemicals, increased chromosome breaks, and elevated γH2A.X foci formation in irradiated cells. Therefore, the global DNA damage in cells lacking FMRP remains to be more broadly tested in other experimental systems using standardized mode of replication stress.

A related key discovery is the correlation between drug-induced DSBs and RLFSs in FX cells, prompting the hypothesis that FMRP is an R-loop processor, and therefore a transcription regulator, which places it functionally upstream of mRNA transport and translation. This new function of FMRP is supported by its ability to suppress RLFS-induced recombination during programmed replication-transcription conflict, that is dependent on its KH2 RNA-binding domain. It is further strengthened by FMRP binding to chromatin sites adjacent to RLFS, which in turn associates with DSBs. It is also consistent with the known affinity of FMRP for, in descending order of degree, RNA, ssDNA and dsDNA (Ashley et al., 1993), three substrates which are all present in an R-loop. Lending further support to our hypothesis we detected co-immunoprecipitation between FMRP and DHX9, a known R-loop regulator. Finally, FMRP was reported to interact with the THO/TREX complex, a mRNP transporter known to be involved in R-loop prevention (Dominguez-Sanchez et al., 2011), through affinity purification (Hein et al., 2015). This new function of FMRP has profound implications for FXS etiology as it adds another layer of complexity to the impact of FMRP deficiency. It also suggests a potential therapeutic intervention by targeting co-transcriptional DNA:RNA hybrids in FX cells.

### A potential mechanism linking dysregulated protein synthesis to genome instability in FX cells

FMRP is predominantly localized to the nuclear periphery and in the cytoplasm in the absence of DNA damage. Replication stress induces co-localization of FMRP and RNA:DNA hybrids towards the center of the nucleus in the lymphoblasts. Fibroblast nuclei staining further suggests RNA:DNA hybrids localizing to the nucleous in FX cells. These results are consistent with the observed correlation between DSBs and RLFSs, preferentially occurring on rDNA-bearing chromosomes, which are resident of the nucleolus. Are rDNA-bearing chromosomes prone to DSBs simply by virtue of being proximal to the nucleolus as a locale, or is there an underlying cause for the them to generate RLFS-associated DSBs? We favor the latter explanation. While the 45S rDNA array residing on the short arms of five acrocentric chromosomes (13, 14, 15, 21, and 22) defines the nucleolus, the 5S rDNA array is a resident of chromosome 1 and the 5S and 45S rDNA arrays are not in close proximity spatially in human lymphoblastoid cells (Yu and Lemos, 2016). This suggests that DSB-RLFS association on these chromosomes is not mediated by proximity to the nucleolus *per se*. Instead we reason that the act of rRNA transcription subjects these chromosomes to increased R-loop formation and chromosome breakage, outside the rDNA arrays. FMRP deficiency leads to elevated level of protein translation (Darnell et al., 2011), which would be reliant on increased rate of ribosome production. Consequently, FX cells must sustain high levels of ribosome production and rRNA transcription, and thus relay the stress, likely torsional in nature, from the rDNA loci intra-chromosomally onto the remainder of the chromosome. Torsional stress of the chromosome, when combined with replication stress, would then induce heightened replication-transcription conflicts and chromosome breakage.

### Novel cellular pathways linked to FXS through genome instability

Neuronal development genes appeared susceptible to strand breakage in FX cells even without drug treatment. Additionally, genes containing overlapping RLFSs and drug-induced DSBs in FX cells are enriched in flavonoid glucuronidation. Moreover, genes containing FMRP-binding sites are also enriched in flavonoid metabolism. These results led us to hypothesize that FX cells, when under stress, are defective in glucuronidation of xenobiotics. Glucuronidated flavonoids have a reported protective role towards a range of neurological disorders (Docampo et al., 2017). Conversely, decreased glucuronidation of xenobiotics such as bisphenol A has been observed in patients with Parkinson’s Disease (Landolfi et al., 2017). Finally, metabolic profiling of *FMR1* premutation carriers who have intermediate (55-200 copies) CGG repeat expansion but nevertheless manifest FMRP deficiency, showed elevated levels of glucuronic acid (Giulivi et al., 2016). We found FX-specific DSBs in genes coding for the most important enzymes in phase I (cytochrome P450 enzymes, specifically CYP2C9 and CYP2C19) and phase II (UGTs, specifically the UGT1A subfamily isoforms) xenobiotics metabolic pathways. We further demonstrated that both UGT1 and CYP2C9 expression levels are reduced in FX cells. The UGT1A subfamily enzymes glucuronidate bilirubin, xenobiotic phenols, and a wide range of psychotropic drugs (de Leon, 2003). Bilirubin glucuronidation is catalyzed by UGT1A1, the single enzyme that may be spared by DSB formation at the *UGT1* locus. Consistently, we have not come across any report of hyperbilirubinemia in FXS patients. Therefore we suggest that FX individuals are defective in metabolizing xenobiotics and psychotropic drugs, which can lead to neurotoxicity and further perils of the neurological functions.

In summary, our study led us to present a model (Fig. 6) where FMRP associates with its gene substrates co-transcriptionally. Upon drug-induced double insults of replication stress and unscheduled gene expression, FMRP increases its nuclear presence to prevent R-loop formation and chromosome breakage during heightened replication-transcription conflicts. We note that FX cells also produce spontaneous DSBs at a higher level than NM cells, and these spontaneous DSBs are not correlated with RLFSs. This suggests that FMRP has additional protective role(s) towards the genome without external replication stress. Recent studies have shown that FMRP deficiency causes imbalance of epigenetic modifications due to unregulated protein synthesis (Korb et al., 2017; Li et al., 2018). It is plausible that the spontaneous chromosome breakage in FX cells is a result of altered histone modifications predisposing specific regions of the chromatin to breakage. Together with our discovery that FMRP directly interacts with the chromatin, these attributes make FMRP a novel mediator of transcription and replication whose function is to prevent R-loop accumulation and ensure genome integrity, thereby maintaining normal synaptic plasticity in neuronal cells. Finally, our study marks a technological advance in mapping chromosome breaks by Break-seq, previously developed for the model organism *Saccharomyces cerevisiae* (Hoffman et al., 2015). To this date, Break-seq has shown efficacy in multiple mammalian cell systems including suspension cell culture (this study), adherent cell culture and 3D organoids (unpublished). Thus, we believe Break-seq holds tremendous potential as a powerful tool for genome discoveries.

**Figure 6.**
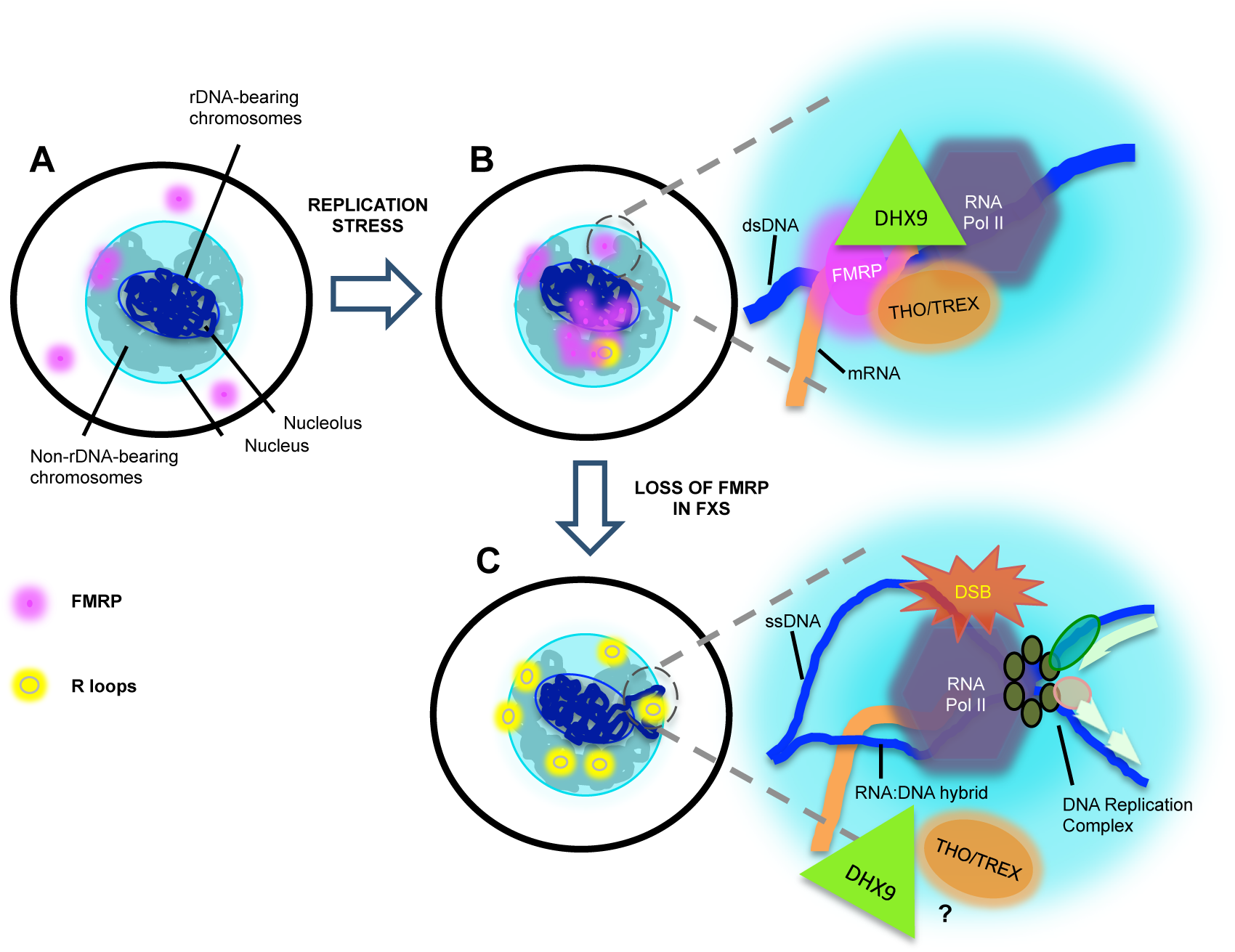
Proposed model of a genome maintenance function of FMRP in the nucleus. **(A)** Illustration of a normal lymphoblastoid cell without any treatment, showing FMRP in the cytoplasm and in the nuclear periphery possibly engaged in mRNA transport. **(B)** Under replication stress, FMRP localizes to the center of the nucleus at R-loop formation sites in genes induced by DMSO and APH. At the junction of convergent replication and transcription, FMRP in conjunction with DHX9 and possibly the THO/TREX complex is involved in R-loop removal and avoidance of a deleterious collision (inset). dsDNA, double-stranded DNA. **(C)** In FX cells increased protein synthesis rate demands high level rRNA production on the rDNA-bearing chromosomes, which in turn causes increased level of (RNA Pol II) transcription elsewhere on these chromosomes, represented by the chromosome loops tethered to the nuclear pores for active transcription. Absence of FMRP permits stable R-loop formation and DSBs upon collision of replication and transcription (inset). The involvement of DHX9 and the THO/TREX complex in this context remains to be determined. ssDNA, single-stranded DNA.

## DATA ACCESS

All primary sequencing data described in this manuscript have been deposited in the NCBI Gene Expression Omnibus (GEO; http://www.ncbi.nlm.nig.gov/geo/) under the accession number GSE124403. They can be viewed through the following link provided by GEO while they remain in the private status until publication: http://www.ncbi.nlm.nih.gov/geo/query/acc.cgi?token=aritcgkovvknxoz&acc=GSE124403

## ACKNOWLEDGEMENTS

We wish to thank V. Van Steenkist for systems and programming support; Drs. C. Dobkin, B. Howell and A. Aguilera for generous gift of antibodies and plasmids; and Drs. X. Chen, L. Kotula, F. Middleton, P. Kane and T. Wongsurawat for helpful discussions. We also thank the staff at SUNY Upstate Flow Cytometry Core and the University at Buffalo Genomics Core for flow sorting and HiSeq sequencing, respectively. This work was supported by the NIH awards (5R00GM08137805 and 5R01GM11879901) and the Department of Defense CDMRP Discovery award (PR141850) to W.F. and institutional support from A*STAR and SUNY EMPIRE scholar grant to V.A.K.

## AUTHOR CONTRIBUTIONS

A.C., A.M., B.H., E.H., and W.F. conceived the study and designed the experiments. A.C., A.M., E.H., A.B. and W.F. performed the Break-seq experiments. A.C. and B.H. performed the ChIP-seq experiments. A.C., J.L., and H.H. performed and analyzed the immuno-staining experiments. A.C. and A.T. performed the recombination assays. A.C. performed and analyzed the flow cytometry, chromatin fractionation, and ChIP-qPCR studies. P.J., V.K., S.E.H., C.C. and W.F. performed computational analyses. A.C. and W.F. wrote the manuscript with input from all authors;

## MATERIALS AND METHODS

### Cell line growth and drug treatment conditions

Human EBV transformed lymphoblastoid cell lines, GM06990 (control) and GM03200 (Fragile X), and fibroblast cell lines, GM00357 (control) and GM05848 (Fragile X), were purchased from Corielle institute. Lymphoblastoids were grown in RPMI1640 (Corning cell gro), supplemented with GlutaMAX (GIBCO), 15% heat-inactivated FBS (Fetal Bovine Serum, Benchmark), 100 IU/mL penicillin and 100 µg/mL streptomycin (Corning cell gro) at 37°C with 5% CO_2_. Fibroblast cells were cultured in MEM culture media with 15% FBS (Corning), 1X GlutaMAX, 100 IU/mL penicillin and 100 µg/mL streptomycin. Lymphoblast cells were treated, at a density of 0.4-0.5×10^6^ cells/ml, with 0.03 μM, 0.3 μM, or 0.6 μM APH (A. G. Scientific), solvent (DMSO, 0.02%, same as the concentration in the APH-treated samples) only, or nothing, for 24 h before harvest. Fibroblasts were treated at 30-40% confluency with the same drug concentrations.

### Flow cytometry for cell cycle analysis

Approximately 1.5-2×10^6^ cells from the Break-seq experiments were harvested for flow cytometry. Cells suspended in 1 ml of PBS were slowly added to chilled absolute ethanol and stored in −20°C. Fixed cells were pelleted at 250xg for 15 m at room temperature and then rehydrated with 5 ml PBS for 15 m. Cells were again pelleted and resuspended at 0.5×10^6^ cells/ml in propidium iodide solution (40 µg/ml propidium iodide, 100 µg/ml RNase A in PBS) and incubated for 20 m at 37°C. Cells were passed through filter-topped flow tubes (BD Falcon) using a luer-lock syringe and analyzed using Becton Dickinson Fortessa Cell Analyser (BD Biosciences). Data were analyzed by FlowJo.

### Flow cytometry for quantification of cells stained for γH2A.X

Approximately 3×10^6^ cells were treated with APH, DMSO or nothing. For compensation control, an additional 3×10^6^ cells were subjected to 2 flashes of UV irradiation at 20 μJ/cm^2^ and allowed to recover for 4 h before harvest. Cells were treated with 1:500 diluted Zombie Aqua (Violet, Biolegend) to stain dead cells, followed by a wash in FACS buffer (2% FBS in PBS). Cells were then fixed in 500 µl of 4% paraformaldehyde, permeabilized by 500 µl methanol, and stained with 100 µl of a 1:50 dilution of anti-γH2A.X in dilution buffer (0.5% BSA in 1x PBS) for 1 h. Cells were then centrifuged and washed in dilution buffer followed by 1x PBS. Cells were resuspended in FACS buffer and filtered through filter-topped flow tubes (BD falcon) using a luer-lock syringe. Samples were analyzed using Becton Dickinson Fortessa Cell Analyser (BD Biosciences) and data analyzed by FlowJo. Dead cells were removed from the analysis. Stained but untreated NM cells were used to generate a baseline for fluorescence. Cells with DNA damage were gated based on fluorescence intensities (FI) higher than the baseline. Percentage of cells with DNA damage were calculated based on the number of cells above the baseline FI and total live cells.

### Immunocytochemistry and microscopy

*For lymphoblasts:* Approximately 3×10^6^ cells having undergone drug treatment described above were washed twice in PBS before fixing with 500 μl of methanol or 4% paraformaldehyde in microfuge tubes. *For firbroblasts:* Approximately 1×10^5^ cells were plated on poly-D-lysine (Sigma Aldrich)-coated coverslips and cultured for 72 h, followed by drug treatment for 24 h. *For both lymphoblasts and fibroblasts:* Cells were washed with 500 μl 1X PBS twice, fixed with 500 μl 4% paraformaldehyde for 20 m at room temperature, followed by gently washing with 1X PBS three times. Cells were then blocked with 500 μl PBSAT (1% BSA, 0.5% Triton X in 1X PBS), followed by incubation with 100 μl of primary antibody solution for 1 h, washed with PBSAT, and incubation with 100 μl secondary antibody for 1 h. Cells were then washed with PBSAT followed by PBS, and resuspended in mounting media (Prolong Diamond antifade plus DAPI, Invitrogen) before being placed as a drop onto microscope slides. Coverslips were carefully placed on top of the mounted drops and allowed to solidify for 24 h before imaging on Leica STP 800 wide-field fluorescence microscope (for lymphoblasts) or Leica SP8 confocal (for fibroblasts). Antibodies used for immunostaining include the following: primary antibodies (anti-γH2A.X, Cell Signaling, 1:500; anti-FMRP, Cell signaling, 1:200; S9.6, Kerafast, 1:500 for lymphoblasts and 1:120 for fibroblasts; and anti-Lamin A&C, Novus Biologicals, 1:250) and secondary antibodies (Alex fluor 488, 568, and 647, Invitrogen, 1:400).

To quantify γH2A.X staining signals maximum projection of 3D image stacks acquired from thirty-six 2D imaging planes with a step size of 0.2 micron along the z-axis was performed using the MetaMorph software (Molecular Devices). Image stacks were deconvolved using the AutoQuant software. In Fiji, DAPI was used to create region of interest (ROI) of nuclei in γH2A.X channel for individual cells. Maximum intensity projections adjusted for background in Fiji were used to quantify γH2A.X intensities in ROI for approximately 28-35 cells. Representative images adjusted for background and contrast are shown. FIRE LUT was used to show intensities of foci per nuclei in every sample. 3D surface plots were also conducted in Fiji (plot parameters: filled, 0.2 perspective, FIRE LUT, 0.88 scale, 0.87 z-scale, 100% max and 19% min) as a measure of the intensities of foci per nuclei per sample. Statistical analysis was done using GraphPad Prism 7 and values were plotted in Kaleidagraph.

To determine co-localization of FMRP and R-loop in the nucleus, 3D image stacks were acquired from sixty-one 2D imaging planes with a step size of 0.11 micron using Metamorph. For images shown in Fig. 5 a single Z-plane image at approximately the center of the stack was shown for each sample. DAPI was used to create a ROI which was overlayed and colored white to indicate nucleus in all three channels (DAPI, FMRP, S9.6). Images were adjusted for background and contrast and smoothed using a gaussian blur of 0.7 in Fiji. 3D construction of the image stacks were performed in Metamorph with rotation along the X-axis every 10° for FMRP and S9.6 channels, followed by conversion into a movie using Metamorph.

### Single cell gel electrophoresis assay

Neutral comet assay was performed using the TREVIGEN reagent kit (cat# 4250-050-K) according to the manufacturer’s instructions. Comet images were analyzed using the CaspLab software (Konca et al., 2003).

### Break-seq and ChIP-seq

Lymphoblastoids GM03200 and GM06990 were used for Break-seq and ChIP-seq analyses. For Break-seq three independent sets of experiments were performed, wherein Set A and B were technical replicates from the same experiment and Set D and E were biological replicates. Break-seq library construction was performed as previously described (Hoffman et al., 2015) with modifications. ChIP-seq analysis, detailed in the Supplemental Information, was followed by independent validation of target genes by ChIP-qPCR.

### Calculation of replication timing for DSB regions

Replication timing data were derived from Repli-seq data of lymphoblastoid GM06990 cells (accession: ENCSR595CLF) publicly available from ENCODE (https://www.encodeproject.org/replication-timing-series/ENCSR595CLF/). An S50 (0<S50<1) value, defined as the fraction of the S phase at which 50% of the DNA is replicated (50% of the cumulative enrichment), was computed for any 50-kb segment of the genome (Hansen et al., 2010). The cumulative enrichment was calculated for each sliding window of 50 kb at a 1-kb step size by linear interpolation of enrichment values in 6 evenly divided temporal windows of the S phase, as previously described (Chen et al., 2010). If a given 50-kb segment was not significantly enriched in any window in the S phase, no S50 value was attributed (S50=NA). Approximately 5% of the genome fell in this category. The DSB regions were then assigned the same S50 values as that of the 50-kb segment in which they reside. For FX cells DSBs on the Y chromosome were excluded from further analysis due to the lack of replication timing data in the reference genome of GM06990. Finally, the DSBs with assigned replication timing values were further parsed into early (S50<0.5) and late (S50>0.5) replicating domains. The resulting distribution of DSBs in the early and late replicating domains was subjected to a Genomic Association Test (GAT) to determine if the DSBs were enriched in either of the two domains through 1000 randomized simulation (Heger et al., 2013).

### Co-immunoprecipitation (Co-IP)

Approximately 6-7 x10^6^ cells were used for each IP reaction. Cells were resuspended in 1 ml IP lysis buffer [25 mM Tris-HCl pH 7.5 / 150 mM NaCl / 1% NP-40 / 1 mM EDTA / 5% glycerol / Halt protease inhibitor cocktail (Thermo scientific) / Halt phosphatase inhibitor cocktail (Thermo scientific)] and incubated on ice for 1 h. Cell lysates were centrifuged at 10,000 rpm for 10 min. Protein concentration in the supernatant was determined using Pierce protein assay reagent (Thermo Scientific). 50 µl of Dynabeads protein G (Invitrogen) per reaction were incubated with 200 μl antibody binding buffer [1X PBS/ 0.02% Tween 20] and 5 μg of anti-FMRP (Covance), or anti-DHX9 (Santa Cruz Biotechnology), or IgG (Biolegend) in a rotator for 10 m at room temperature. The immuno-complex was rinsed with 200 μl antibody binding buffer at room temperature, followed by incubation with 500 μg of cell lysate per reaction at 4°C overnight. After incubation the supernatant was saved as flow-through (FT) and the beads were washed twice with IP lysis buffer without NP-40, saving each wash. 50 μl 2X Laemmli buffer was added to the beads and boiled for elution, before analysis on 8% SDS-PAGE gels and western blotting using anti-FMRP (Cell signaling, 1:500), anti-GAPDH (Thermo scientific, 1:4000) or anti-DHX9 (Santa Cruz Biotechnology, 1:500).

### Subcellular fractionation

GM06990 and GM03200 cells were grown to a density of 0.4-0.5×10^6^ cells/ml with >90% viability. Cells were treated for 24 h with aphidicolin, DMSO or nothing. Samples were collected as aliquots of approximately 5×10^6^ cells, washed twice with PBS, then frozen for storage. Each thawed aliquot of cells was resuspended in 500 µl Farham’s lysis buffer without NP-40 [5 mM PIPES pH 8.0 / 85 mM KCl / Halt protease inhibitor cocktail] and incubated on ice for 2 m. Fifty-µl of the cell lysate thus prepared was collected as a whole cell extract control and the remaining lysate was spun at 1300xg for 4 m to pellet nuclei. The supernatant served as the crude cytoplasmic fraction. The nuclear pellet was resuspended in 150 µl Farham’s lysis buffer and incubated for 20-30 m at 4°C and served as the nuclear fraction. Equal volume of 2X Laemmli buffer were added and samples were boiled and later sonicated. Approximately 3×10^5^ cell equivalent per fraction was used for electrophoresis on a 12% SDS-PAGE gel, followed by western analysis. Densitometry of autoradiogram was done using ImageJ (https://imagej.nih.gov/ij/) to calculate the percentages of FMRP in the nuclear and cytoplasmic fractions.

### Western blot

For protein expression in human cell lines, whole cell lysates were prepared in lysis buffer [50 mM Tris-HCl pH 7.5 / 0.5 M NaCl / 10 mM MgCl_2_ / 1% NP-40 / Halt protease inhibitor cocktail / Halt phosphatase inhibitor cocktail] and at least 20 μg of proteins were analyzed by 10% SDS-PAGE before western blotting. The following antibodies were used: anti-FMRP (Covance, 1:1000), anti-UGT1A (Santa Cruz Biotechnology, 1:250), anti-CYP2C9 (Invitrogen, 1:1000), anti-Histone H3 (Cell Signaling, 1:500) and anti-GAPDH (Thermo scientific, 1:2000). For yeast whole cell extracts, a single colony was inoculated in 10 ml SC-HIS-URA and grown overnight. Cells were centrifuged at 3000 rpm for 5 m, frozen and stored at −80°C until further use. Frozen cell pellets were thawed in 250 µl TBS [50 mM Tris pH 7.5 / 100 mM NaCl / Halt protease inhibitor cocktail (Thermo scientific)]. Cell suspension was vortexed at 4°C for 15 m after the addition of 200 μl sterile glass beads, followed by centrifugation at 13500 rpm at 4 °C for 10 m. The supernatant was retained and protein concentration was determined by Bradford assay (BioRad). Approximately 25μg of proteins were analyzed on 10% SDS-PAGE followed by western blots using monoclonal anti-FLAG-M2-Peroxidase (HRP) (Sigma Aldrich, 1:2000) and anti-β actin (1:4000).

### Yeast strains and plasmids

Yeast strains used in this study were BY4741 (*MATa his3Δ1 leu2Δ0 ura3Δ0 met15Δ*) and its isogenic derivative *rnh1Δ* (*MATa ura3Δ0 leu2Δ0 his3Δ1 met15Δ0 rnh1Δ::KAN*, EUROSCARF collection (Entian et al., 1999)). Yeast cells were either grown in YEPD or synthetic complete (SC) media with specific amino acids omitted as indicated. All yeast strains were grown at 30°C with horizontal shaking for liquid cultures. Yeast centromeric plasmids pARS-GLB-OUT (OUT) and pARS-GLB-IN (IN) containing GAL-OUT/IN recombination constructs were provided by and described previously (Prado and Aguilera, 2005). Specifically these plasmids were designed with the *leu2Δ3*’::*leu2Δ5*’ direct-repeat recombination system under the *GAL1* promoter. The plasmid also contains *ARSH4, URA3, CEN6* and the 83 bp (C-A1-3)_n_ telomeric sequences from pRS304 lacking the *EcoR*I site at the polylinker. Human gene sequences for recombination assay (see below) were cloned between the direct-repeat recombination system (*leu2Δ3’:sequence: leu2Δ5’*) using *Bgl*II. The following primers containing *Bgl*II site in the forward and *BamH*I site in the reverse were used for the described sequences: a) control-1_**F**-5’-TCagatctTCAGGCTGCACATTCTTTTC-3’ and control-1_**R**-5’-CTCggatccTGCTTTCACTGCAGTTCC-3’; b) control-2_**F**-5’-CTCagatctTGATAATTACAAGGTACACGTTATTGC-3’ and control-2_**R**-5’-CTCggatccTTGGTTAGGATAATAAGCACTATGG-3’; c) RLFS-1_**F**-5’-CTCagatctGTAGACGCCTCACCTTCTGC-3’ and RLFS-1_**R**-5’-CTCggatccTGCGGGTGTAAACACTGAAA-3’; d) RLFS-2_**F**-5’-CTCagatctCATAACTAAGCACTGTATGCC-3’ and RLFS-2_**R**-5’-CTCggatccCCTAGGGACAAGGGGAGGTA-3’.

The above sequences were PCR amplified from human genomic DNA (http://www.fenglab-genomestabilityresearch.org/p/isolation-of-genomic-dna-from-mammalian.html). Sequences were inserted in two orientations due to compatibility of ends generated by *Bgl*II and *BamH*I and both the orientations were used to measure recombination frequencies for all sequences. The sequences were inserted in both the IN and the OUT constructs. pFRT-TODestFLAGHAhFMRPiso1 plasmid (Addgene) was used to subclone an *Spe*I/*Bcl*I-digested CMV-FMRPiso1 fragment into pRS316 at the *Xba*I and *BamH*I cloning sites. The resulting construct, pRS316-CMV-FMRPiso1, was then digested with *Not*I and *EcoR*I to obtain the CMV-FMRPiso1 fragment. The fragment was subcloned into pRS313 digested with *Not*I and *EcoR*I producing the final construct pRS313-CMV-FMRPiso1. pRS313-CMV-FMRPiso1I304N was generated using the same procedure from the pFRT-TODestFLAGHAhFMRPiso1I304N plasmid (Addgene).

### Recombination frequency assay

The IN and OUT plasmids were first transformed in BY4741 or *rnh1Δ* and selected in SC without uracil in 2% glucose. The fluctuation assay was performed as previously described with modifications (Prado and Aguilera, 2005). Briefly, selected transformants were streaked onto SC-URA+2% Glucose and SC-URA+3% Galactose. Plates were incubated for 4 days at 30°C to suppress or induce transcription through the *GAL1* promoter respectively. Six single colonies for every sample were re-suspended in 1 ml –N media (1.61 g/l YNB without (NH4)_2_SO_4_ or amino acids, 94 mM succinic acid and 167 mM NaOH) and sonicated. Serial dilutions were prepared for each of the six colonies per sample: 1:15, 1:150 and 1:1500 in a 96-well plate. 100 µl of diluted samples were plated in SC-URA+3% Galactose for calculation of totals. For calculation of recombinants; 100 µl from undiluted was plated onto SC-LEU-URA+3% Galactose. Plates were incubated for 3 days at 30°C, and colonies were counted to calculate recombination frequency as follows:

> no. of recombined colonies/(total no. of cells plated*dilution factor)*10^4^

Recombination frequency was calculated for each of the six colonies per sample and the median value was used as the recombination frequency of a sample. Three independent experiments were conducted for each construct and treatment (glucose and galactose) and standard deviations were calculated for graphical representation and to estimate error.

For experiments with FMRP expression the IN and OUT plasmids with or without RLFS were co-transformed with pRS313-CMV-FMRPiso1, pRS313-CMV-FMRPiso1I304N or pRS313 into BY4741 and selected in SC-URA-HIS+2% glucose. Recombination frequency assay was conducted as described above with the totals plated in SC-URA-HIS+3% galactose and the recombinants were plated in SC-LEU-URA-HIS+3% galactose.

### Gene ontology analysis and identification of pathways that are potentially altered by treatment

Gene ontology analyses are performed via DAVID Bioinformatics tools (https://david.abcc.ncifcrf.gov), DiffBind, or WebGestalt (http://webgestalt.org).

### Genomic association tests for correlation between DSBs and CFS cores and the “APH.breakome”

The DSB regions with assigned replication timing indices (early vs. late, see Methods in main text) were compared to previously published finely mapped CFS core sequences (Savelyeva and Brueckner, 2014) and the “APH.breakome” (Crosetto et al., 2013), using the Genomic Association Tester (GAT) software (Heger et al., 2013). In all tests the DSBs were set as *segments* and the other datasets as *annotation*, with the genomic regions previously assigned with S50 values as *workspace* (*i.e.,* excluding those regions with S50 value of “NA”) and default parameter for sampling rounds (--num-samples=1000).

### Statistical analysis

Two-way ANOVA test followed by Tukey’s multiple testing for all pair-wise comparisons was performed for all experiments unless otherwise noted. Annotation for P values in figure legends regardless of statistical test type are: *, p<0.05; **, p<0.01; ***, p<0.001; ****, p<0.0001. Error bars denote standard deviation unless otherwise noted.

