## Supplemental Information for "Fragile X Mental Retardation Protein regulates R-loop formation and prevents global chromosome fragility"

#### SUPPLEMENTAL METHODS

##### Break-seq

***Break-seq library construction.*** Break-seq procedures were as described previously with modifications (Hoffman et al., 2015). 5x$10^6$ cells were embedded into 0.5% Incert low-melting point agarose in PBS and cast into plugs. The agarose plugs were then incubated at 50ºC overnight in 6 ml of lysis buffer (0.5 M EDTA, 1% Sarkosyl, 200 µg/ml Proteinase K). The DNA in the agarose plugs was then end-labeled in-gel using the End-It Kit (Epicentre) with biotinylated dNTP mix (1 mM dTTP, dCTP, dGTP, 0.84 mM dATP, 0.16 mM Biotin-14-dATP). Plugs were then treated with β-Agarase (NEB) to digest agarose and release DNA. DNA sample was then sonicated using a Covaris M220 using the snap-cap DNA 300 bp shearing protocol. DNA was then processed using a PCR Cleanup Kit (Qiagen) and run on agarose gel to verify the fragmentation pattern of DNA and quantified on a Nanodrop. 10-11 µg of DNA was then end repaired (Epicentre) and purified by the PCR Cleanup Kit (Qiagen). The DNA was then A-tailed by A-tail Kit (NEB) or Klenow exo- (NEB E6054A) and purified by PCR Clean-up Kit (Qiagen), followed by quantification on a Nanodrop. M270 Dynabeads (Life Technologies) were used to purify biotinylated DNA. The amount of DNA bound to beads was calculated by measuring the quantity of DNA in the flow through. DNA-bound beads were then resuspended in ligation mix containing Illumina adaptors (50 µM adaptor-1, 50 µM adaptor-2, 1x T4 ligase buffer, 3 µl T4 DNA ligase) and incubated overnight at room temperature on a roller. 400 ng of DNA bound to beads was used for PCR amplification using KAPA Hotstart Ready Mix (KAPA). Each sample was given a specific index primer for multiplexing. PCR product was then run on agarose gel to verify amplification and quantity. AMPure beads (Agencourt) were used to remove free adaptors and the final product was analyzed on agarose gel. Break-seq libraries were sequenced on Illumina Hi-Seq 2500 with 100 or 150 bp paired-end reads, followed by Break-seq data analysis. Adaptor sequences and index primer sequences were previously described (Hoffman et al., 2015).

***Break-seq DSB peak identification***. Raw sequence reads were obtained from Illumina Hi-seq 2500 and then aligned to the UCSC human genome assembly, GRCh37/hg19 (http://hgdownload.soe.ucsc.edu/goldenPath/hg19/bigZips/), using Bowtie 2 (http://bowtie-bio.sourceforge.net/bowtie2/index.shtml) in the "--*local"* mode. The PCR duplicate reads were removed using Picard MarkDuplicates (http://broadinstitute.github.io/picard). The non-redundant mapped sequence reads were sorted and then converted to BAM files using SAMtools (Li et al., 2009) (http://samtools.sourceforge.net/) and subjected to subsequent processing with Model-based Analysis for ChIP-seq (MACS version 2.1.1, https://pypi.python.org/pypi/MACS2) using two-sample analysis between break-seq samples (treatment) and whole genome sequencing data (control) using the *callpeak* function in MACS2 with a p value <1e-5. For identification of *Pac*I-digested breaks, DSB peaks with perfect match to *Pac*I motif (TTAATTAA) were mapped onto the hg19 reference genome with Bowtie. *IntersectBED* function from BEDtools (Quinlan and Hall, 2010) was then used to find overlap between *Pac*I motif sites and peaks identified through MACS2. These overlapping peaks were considered *Pac*I sites found in the Break-seq sample.

***Random permutation tests for identification of PacI sites***. The *shuffleBed* function in BEDtools was used to randomly permute the genomic locations of DSBs identified as *Pac*I sites with default parameters to generate randome genomic locations as a null distribution, preserving the size of DSBs and number of DSBs per chromosome. The fraction of sequences containing *Pac*I motif was calculated. One thousand iterations of this process was performed. The distribution of the *Pac*I-positive fractions was then compared to that from the experimental dataset and One Sample Student's t-test was performed.

***Break-seq library complexity calculation and identification of consensus DSB peaks in replicate experiments.*** All biological replicates for each sample (strain/treatment combination) were pooled for assessement of library complexity by preseq (Daley and Smith, 2013). All 23 Break-seq peak files were analyzed in DiffBind (Ross-Innes et al., 2012) for consensus DSB peak identification. Consensus DSB peaks were defined as those that appear in at least two replicate experiments, regardless of the total number of replicates, for each sample (cell line/treatment). Aphidicolin-treated samples (0.03 µM and 0.3 µM) were combined to form a composite APH-treated sample, for both NM and FX cells, and consensus DSB peaks were then extracted similarly as described.

***Correlation between DSBs and other genomic features.*** The association between DSBs and other genomic features including RLFSs and DRIP-seq signals was determined using the *bedtools annotate* function. Multiple data sets of DRIP-seq were concatenated (*cat GSE70189_NT2_DRIPc_peaks GSM1720615_NT2_DRIP_1_peaks GSM1720616_NT2_DRIP_2_peaks GSM1720617_NT2_DRIP_RNaseA_peaks GSM1720618_NT2_DRIP_RNaseH_peaks GSM1720619_K562_DRIP_peaks.clip > composite.DRIP*), sorted (*sort -k1,1 -k2,2n composite.DRIP > composite.DRIP.sorted*) and then merged into a composite data set using the *bedtools merge* function (*bedtools merge –i composite.DRIP.sorted > composite.DRIP.sorted.merged*). The significance of the association or p value was calculated using the fisher exact test (*fisher*) in BEDtools. Relative distance between DSBs and RLFSs was calculated using *bedtools reldist* function.

**ChIP-seq**

***Cell collection.*** GM06990 and GM03200 cells were grown to log phase with a viability >90%. Cell fixation, harvest and IP were conducted according to Richard Myers lab ChIP-seq protocol. Briefly, cells were fixed with 1% formaldehyde for 10 m at room temperature and then blocked with 0.125 M glycine. Cells were then washed twice with cold PBS. Cells were collected at $2x10^7$ cells per ChIP reaction, snap frozen in liquid nitrogen and stored at -80ºC untill further use.

***Immunoprecipitation.*** Cells were thawed on ice with 1 ml Farham's lysis buffer (5 mM PIPES pH 8.0, 85 mM KCl, 0.5% NP-40, Halt protease inhibitor cocktail (Thermo Scientific)) for 5 m. Nuclei were prepared by centrifuging the lysate at 2000 rpm for 5 m. The nuclear pellet was resuspended in 300 µl RIPA (1x PBS, 1% NP-40, 0.5% sodium deoxycholate, 0.1% SDS, Halt protease inhibitor cocktail). Samples were sonicated in a Bioruptor (Diagenode) at high setting for a total time of 40 m with 30 s ON and 30 s OFF at 4°C. The sonicated mixture was centrifuged at 13,500 rpm for 15 m at 4°C and the volume of supernatant was adjusted to 1 ml with RIPA buffer. One hundred µl of this mixture was set aside as "input" and the remaining nuclear fraction was used for immunoprecipitation. Both aliquots were snap frozen in liquid nitrogen and stored at -80°C until ready for immunoprecipation. Ten µg of monoclonal anti-FMRP antibody (Covance, now Biolegend, Cat#MMS-5231) was conjugate with M280 Dynabeads sheep anti-mouse IgG (Life Technologies). The thriced washed beads (with

PBS/BSA) were incubated with 900 μl sonicated nuclear fraction overnight at 4°C with agitation. The beads were then washed 5 times with cold LiCl wash buffer (100 mM Tris pH7.5, 500 mM LiCl, 1% NP-40, 1% sodium deoxycholate), followed by a single wash in TE (10 mM Tris-HCl pH7.5, 0.1 mM EDTA). The immunoprecipitates were eluted with 200 μl of IP elution buffer (1% SDS, 0.1 M $NaHCO_3$) at 65°C for 2 h with vortexing every 30 m. The 100 μl "input" was thawed and together with the eluted immunoprecipitates were reverse-crosslinked at 65°C overnight. The samples were then used for ChIP-seq library construction.

***Library preparation.*** DNA was purified from all samples (input and immunoprecipitates described above) using Qiagen PCR clean-up kit and then end-repaired in a 100 μl reaction (1x End-repair buffer, 250 μM dNTP mix, 1 mM ATP, and 3 μl End-It Enzyme (Epicentre)) for 45 m at room temperature. Labeled DNA was purified using Qiagen PCR clean-up kit, A-tailed using Klenow(exo-), purified again with Qiagen PCR clean-up kit, and ligated to Ilumina adaptors using the same conditions as described for Break-seq. Ligated DNA was purified with Qiagen PCR clean-up kit and amplified by PCR using index primers (98°C, 5 m; 18 cycles of 98°C, 20 s; 65°C, 15 s; 72°C, 1 m; followed by 1 cycle of 72°C, 5 m). The PCR products were purified by AmPure beads (AgenCourt) to remove free adaptors.

***ChIP-seq data analysis.*** Raw sequence reads were aligned to hg19 by Bowtie2 the same as for Break-seq data. The non-redundant mapped sequence reads were used for peak calling by Model-based Analysis for ChIP-seq (MACS version 1.4.1). Identification of FMRP-binding sites was done using two-sample analysis between ChIP samples from GM06990 (treatment) and GM03200 (control) with the *callpeak* function and applying a p value < 1E-05.

***ChIP-qPCR.*** ChIP was conducted as before, we additionally used IgG along with FMRP antibody as a control in GM06990 cells only. After immunoprecipitation and DNA isolation, the ChIP'ed DNA and the input DNA was diluted 1:10 in water. The PCR reaction was carried out in 10 μl volume. 5 μl of iTaq Universal SYBR Supermix (BioRad), 300 nm of forward and reverse primers (Supplemental Table S6), 3 μl DNA and water made up the reaction mixture. CFX connect real time system (BioRad) was used for qPCR with the following thermal cycling protocol; 95ºC for 30 s, cycle: 95ºC for 10 s and 58ºC for 30 s. This was followed by melt curve analysis from 65ºC to 95ºC by an increment of 0.5ºC for 5 s. 45 cycles were used for every reaction.

#### SUPPLEMENTAL RESULTS AND DISCUSSION

##### Break-seq proof-of-principle experiments and analysis pipeline.

As a proof-of-principle we mapped DSBs produced by *in vitro Pac*I-digestion of DNA from FX cells treated with DMSO or 0.03 µM APH (Supplemental Fig. S3A). More than 96% of DSBs mapped in these two samples correspond to a known *Pac*I site ($p < 2.2e-16$ in random permutation tests with 1000 iterations, see Methods), with 86% concordance between them (Supplemental Table S1). Of the 151,583 *Pac*I sites in the human hg19 genome, 84,672 (56%) and 89,199 (59%) were mapped in the DMSO and APH samples, respectively. These results provided a benchmark for Break-seq with > 97% specificity and > 56% sensitivity (the in-gel digestion efficiency of *Pac*I was ~70% based on Southern blot analysis (data not shown), suggesting a true Break-seq sensitivity of ~80%) . Break-seq library qualities were assessed for read classification (Supplemental Fig. S3B&C) and library complexity (Supplemental Fig. S3D). All Break-seq libraries did not appear to be saturated, but at the current sequencing depth recurrent DSBs were identified. For each strain/treatment combination, *e.g.*, "FX_0.03 µM APH", *consensus DSBs* from at least two replicate experiments, regardless of the total number of replicates, were derived (Supplemental Table S2). The DSBs from 0.03 µM and 0.3 µM APH-treated samples were further pooled into a composite dataset of "APH-treated DSBs", for each cell line, followed by comparison with the DMSO-treated control to identify DSBs shared by DMSO- and APH-treatment as well as those specific to each treatment (Supplemental Fig. S4A&B, Supplemental Table S2). The concordance of DSB peaks between FX and NM cells was determined (Supplemental Fig. S4C).

##### Comparison between DSBs in this study with previously published CFS cores and the "APH.breakome".

APH-induced chromosome breakage defines common fragile sites (CFSs) (Glover et al., 1984). CFSs are also characterized by late replication timing (Hellman et al., 2000; Le Beau et al., 1998; Palakodeti et al., 2004; Wang et al., 1999). We systematically compared the APH-induced DSBs mapped in our study to fourteen reported CFS core sequences (Savelyeva and Brueckner, 2014) and a list of DSBs mapped by a genome-wide technique named BLESS in APH-treated HeLa cells (henceforth "APH.breakome" as named by Crosetto et al) (Crosetto et al., 2013). Indeed, we observed significant correlation between replication stress-induced DSBs in NM and FX cells with the "APH.breakome" ($p < 0.001$, Supplemental Fig. S9A&B). Moreover, the concordant DSBs (those that were shared between our data and the "APH.breakome") were associated with the late replicating regions for both NM and FX cells. This is consistent with the notion that delayed replication timing of intrinsically difficult-to-replicate sequences are prone to DSBs. In addition, the stress-induced DSBs in FX cells were also associated with early replicating regions (Supplemental Fig. S9C&D), which tend to be transcriptionally active and are prone to replication-transcription conflict. However, we did not find significant ($p < 0.001$) correlation between DSBs in any sample with the CFS cores (data not shown). Closer scrutiny of the experimental conditions in CFS studies allowed us to conclude that this apparent discrepancy stemmed from the differential usage of organic solvent for APH, *i.e.*, ethanol vs. DMSO.

Since the first documented usage of ethanol and DMSO as the solvent for APH and the induction of CFSs (Glover et al., 1984), different laboratories have taken to use either solvent for their studies. Direct comparison between these two solvents in CFS induction has only been documented—to the best of our knowledge—in a single study using two subjects (Kuwano and Kajii, 1987). This study demonstrated that increasing concentrations of ethanol, but not DMSO, synergistically increased APH-induced CFSs. It also showed that ethanol treatment alone induced CFS formation at a frequency of 2-9% when administered at a range between 0.02% and 1%. Unfortunately, the effect of DMSO alone on CFS induction was not measured. Our study suggests that DMSO, at least when administered at 0.02%, enhances DSB formation (see more below) compared to untreated cells. Nevertheless, the study by Kuwano and Kajii suggested that ethanol and DMSO have different impact on CFS formation, particularly in the context of APH treatment. Among the studies from which the CFS core sequences were derived and compiled by Savelyeva and Brueckner (Savelyeva and Brueckner, 2014) (Table 1 therein), all but one study used ethanol as the solvent. The study by Zimonjic et al (reference 51 therein) used either ethanol or DMSO to map the FRA3B site and the results were an undifferentiated mixture. In contrast, our study as well as the BLESS study (Crosetto et al., 2013) used DMSO as the solvent for APH. Therefore, it appears that CFS formation is a product of both APH treatment and other undefined cellular effects by ethanol or DMSO, rendering comparison between studies that employ differential usage of these solvents rather tenuous.

Our study revealed that DMSO sensitizes RLFS regions for DSBs in both NM and FX cells (Table 1). DMSO is one of the most common solvents for organic compounds and facilitates the delivery of drugs across cellular membranes. It is also known as an antioxidant with a protective role for human tissues by interacting with the hydroxyl group on various substances. Its protective role is exemplified in its ability to reduce the damaging effect on DNA molecules by radiation. However, depending on the concentration and cellular context DMSO can function as an antioxidant or a pro-oxidant (Kang et al., 2017; Liu et al., 2001; Perez-Pasten et al., 2006; Sadowska-Bartosz et al., 2013). Moreover, its potential genome-damaging effect has not yet been evaluated in mammalian cells to the best of our knowledge. Fortuitously we discovered that 0.02% DMSO (the concentration at which we used to dissolve APH) can cause chromosome breakage in human lymphoblasts through an unknown mechanism. We assume this genotoxic effect is not tissue-specific. Thus, it is imperative to understand the full cellular and genomic impact by DMSO to inform therapeutic treatment involving DMSO as a solvent.

#### SUPPLEMENTAL FIGURE LEGENDS

**Figure S1. Validation of CGG repeat expansion at *FMR1* locus and the absence of FMRP expression in the Fragile X cell lines used in this study.** (**A**) The restriction map of the 5.2 kb *Eco*RI fragment in the *FMR1* locus and probe position used for Southern blots are shown. The probe was synthesized by PCR amplification from genomic DNA isolated from GM06990 using the following primers: FMR1-F, 5'-TGGCTTCTCTTTTCCGGTCT-3'; FMR1-R, 5'-GGGTTACCTTTTGCCTCCTT -3'. (**B**) Southern blots of *Eco*RI-digested genomic DNA from the indicated cell lines. A band corresponding to 5.2 kb (containing 30-50 CGG repeats) was seen in the normal cell lines. In contrast, two bands corresponding to ~7.0 kb and ~7.4 kb were identified in the FX lymphoblastoid and fibroblast lines, which correspond to ~590 and 730 CGG repeat expansion, respectively. (**C**) Western blots confirming the absence of FMRP expression in both lymphoblastoid and fibroblast lines of Fragile X cells.

**Figure S2. Fragile X cells show increased DNA damage. (A)** Box plots for γ-H2A.X immunofluorescence staining of cells under the indicated conditions. Plotted are mean intensity values of fluorescence per nucleus in three independent experiments ($n \geq 28$ nuclei counted for each experiment). One-way ANOVA test followed by Tukey's multiple comparison test were performed. These data were used to calculate the ratios of fluorescence intensities in (B). (**B**) The summary of these experiments as ratios of each sample to the "NM-Untreated" control. Error bars indicate standard errors. (**C**) Representative immunofluorescence microscopy images for γ-H2A.X staining of normal (NM) or Fragile X (FX) cells in vehicle control, dimethyl sulfoxide (DMSO) or aphidicolin (APH) at the indicated concentrations.

**Figure S3. Break-seq data quality check.** Break-seq sample read distribution of a *Pac*I site in the "proof-of-principle" experiment (**A**). (**B&C**) Break-seq library quality assessed by read classifications. "R1" and "R2", Read 1 an Read 2, respectively. Samples with the same treatment were merged to assess overall sequencing depth in (**C**). (**D**) Break-seq library complexity of FX and NM cells ("obs" and "exp" stand for observed and expected library complexities, respectively). Libraries from biological replicates for each treatment condition were merged for complexity measure.

**Figure S4. Break-seq data analysis pipeline.** (**A**) After Bowtie2 mapping, Break-seq data ("DMSO", "0.3 μM APH", and "0.03 μM APH") were normalized for copy number variation by whole genome DNA sequencing ("Total DNA"), for NM and FX cells, respectively (shown as an example for FX cells), during the peak calling step in MACS2. DSB peaks found in at least two replicate experiments for each strain/treatment combination were identified as "Consensus peaks" by DiffBind. Peaks from different APH treatments (0.03 μM and 0.3 μM) were then pooled into a single set of "Consensus peaks in APH", in contrast to the "Consensus peaks in DMSO" set. (**B**) The consensus peaks for each strain/treatment combination (as indicated) were compared with each other to identify overlaps and condition-specific peaks, ready for further comparison with genomic features such as RLFSs. (**C**) Comparison of DSBs between NM and FX cells for like treatments: Untreated, DMSO-treated, and APH-treated.

**Figure S5**. **Control experiments for RLFS-induced DNA breakage and recombination frequency (RF) in yeast.** (**A**) RF is dependent on transcriptional activation. One-way ANOVA followed by Tukey's multiple testing was performed. (**B**) Insertions of RLFS elements from the human genome can further induce RF, specifically when inserted in the "sense" orientation with respect to transcription. (**C**) The RLFS-induced RF can be further enhanced by deletion of *rnh1*, the gene encoding for RNase H1. Note the change of scale on the Y-axis in (C). (**D**) Cells bearing a second plasmid, pRS313, which appeared to have little effect on RF, serve as control for the experiments in Fig. 3B. (**E**) FMRP and FMRP-I304N show similar expression levels in yeast. Cells transformed with a plasmid with or without an RLFS in the LEU2 gene cassette, together with a plasmid with or without the CMV-driven and FLAG-tagged FMR1 or FMR1-I304N, were analyzed by Western blot using the anti-FLAG antibody. Actin served as a loading control.

**Figure S6. FMRP chromatin-binding site analysis.** (**A**) A monoclonal anti-FMRP antibody specifically immunoprecipitates FMRP from GM06990 lymphoblastoids, as shown by western blots. GAPDH served as control. (**B**) FMRP binding site distribution with respect to genes. (**C**) ChIP-qPCR validation of top FMRP binding substrates revealed by ChIP-seq. Relative enrichment (target/control) was calculated as the ratio of fold enrichment ratio (target antibody vs. IgG) for the target gene to the fold enrichment ratio (target antibody vs. IgG) for a control region, and was plotted on the Y axis. Error bars stand for standard errors in three independent experiments. (**D**) ChIP-seq enrichment ratios for fractions of FMRP chromatin binding sites with (pink) or without (blue) overlapping RLFSs. (**E**) Learning and memory genes enriched for FMRP-binding sites. All except *SLC6A1* and *KRAS* also contain DSBs, albeit not within 1 kb distance to FMRP-binding sites, in FX cells. (**F**) Sixteen previously validated mRNA substrates for FMRP, shown here as FMRP chromatin-binding sites and/or APH-induced DSBs in FX cells.

**Figure S7**. **FX-specific DSB formation in two phase I drug metabolic enzymes, CYP2C9 and CYP2C19.** IGV screenshots for CYP2C9 (**A**) and CYP2C19 (**B**). (**C**) Representative Western blots of CYP2C9 protein expression with GAPDH as control and quantification using ratio of CYP2C9 to GAPDH derived from three biological replicates. Two-way ANOVA followed by Sidak's multiple testing was performed.

**Figure S8. Validation of observations made with FX lymphoblasts using FX fibroblasts (GM05848) and a sex- and age-matched control fibroblast cell line (GM00357).** (**A&B**) Fibroblast cell line of an FX individual shows increased DNA damage compared to a control fibroblast cell line. Two independent experiments were performed and one representative experiment shown here. At least 41 nuclei per sample were analyzed in each experiment. Representative images for γH2A.X staining in FX and NM cells under the indicated treatment (**A**) and quantification of γH2A.X signals per nucleus (**B**) are shown. Two-way ANOVA test followed by Sidak's multiple testing was performed. Scale bar, 10 μm. (**C**) FX cells show increased RNA:DNA hybrid foci formation in the nucleus upon drug treatment detected by S9.6 antibody immunofluorescence. Nuclear boundary is marked by staining for Lamin A&C. Scale bar, 10 μm. (**D**) Western blots demonstrate decreased expression of UGT1A upon replication stress in FX cells. Two experiments were performed and a representative experiment is shown.

**Figure S9. Genomic Association Test (GAT) for correlations between DSBs from the six indicated samples and the "APH breakome" mapped by BLESS.** (**A**) Numbers of DSBs found associated with APH breakome signals in the indicated genomic regions. "Obs", observed number of DSBs; “Exp”, expected number of DSBs. Log2 transformation of the fold enrichment values (ratios of observed to expected number of DSBs) are reported for the whole genome (**B**), the early replicating regions (**C**), and the late replicating regions (**D**). Those samples marked with an asterisk indicate p values ≤ 0.001.

#### SUPPLEMENTAL TABLES

**Table S1.** Proof-of principle Break-seq mapping of *Pac*I sites in DMSO- and (0.03 μM) APH-treated FX cells.

| **Cell line** | **Treatment** | **No. of DSB peaks** | ***Pac*I-positive** | **Concordance between *Pac*I-positives** |
|---|---|---|---|---|
| FX | DMSO | 84458 | 80819 (96%) | 70630 (87%) |
| FX | APH* | 87727 | 84975 (97%) | 70592 (83%) |

**Table S2.** Number of consensus DSB peaks (detected in at least 2 replicates) in all categories defined by strain/treatment/comparison combinations.

| **DSB category**[1] | **Number of DSBs** |
|---|---|
| FX.APH | 18473 |
| FX.APH003 | 16796 |
| FX.APH03 | 2112 |
| FX.DMSO | 9167 |
| FX.NT | 4149 |
| FXdmso.FXaph.overlap | 6177 |
| FXdmso.FXaph.uniquetoFXaph | 12296 |
| FXdmso.FXaph.uniquetoFXdmso | 2984 |
| FXdmso.FXnt.overlap | 322 |
| FXdmso.FXnt.uniquetoFXdmso | 8845 |
| FXdmso.FXnt.uniquetoFXnt | 3827 |
| NM.APH | 7002 |
| NM.APH003 | 3506 |
| NM.APH03 | 2161 |
| NM.DMSO | 3927 |
| NM.NT | 2111 |
| NMdmso.NMaph.overlap | 3209 |
| NMdmso.NMaph.uniquetoNMaph | 3792 |
| NMdmso.NMaph.uniquetoNMdmso | 714 |
| NMdmso.NMnt.overlap | 1651 |
| NMdmso.NMnt.uniquetoNMdmso | 2276 |
| NMdmso.NMnt.uniquetoNMnt | 458 |
| FXdmso.NMdmso.overlap | 644 |
| FXdmso.NMdmso.uniquetoFXdmso | 8523 |
| FXdmso.NMdmso.uniquetoNMdmso | 3283 |
| FXnt.NMnt.overlap | 1753 |
| FXnt.NMnt.uniquetoFXnt | 2396 |
| FXnt.NMnt.uniquetoNMnt | 358 |
| FXaph.NMaph.overlap | 4369 |
| FXaph.NMaph.uniquetoFXaph | 14104 |
| FXaph.NMaph.uniquetoNMaph | 2633 |

[1]Strain=FX, NM; Treatment=NT (untreated), DMSO, APH003 (0.03 μM APH), APH03 (0.3 μM APH), APH (composite of APH003 and APH03); Comparison=overlap (shared by two samples), uniquetoXXxx (unique to the sample indicated).

**Table S3.** Correlation between DSBs in various groups and 169222 computationally predicted RLFSs (Wongsurawat et al., 2012) and 108011 composite DRIP-seq signals merged from all DRIP-seq data sets (NT2 and K562) (Sanz et al., 2016). Note the merged DRIP-seq dataset has 6.6 times the coverage of the RLFSs.

| DSB group | Query (RLFS or DRIP-seq) | Number of DSBs | Number of overlaps | Number of possible intervals | P value Left | P value Right | P value Two-tail | Ratio |
|---|---|---|---|---|---|---|---|---|
| FX.NT | RLFS | 4149 | 133 | 2000047 | 2.34E-43 | 1 | 4.37E-43 | 0.358 |
| FX.DMSO | RLFS | 9167 | 1294 | 1702520 | 1 | 3.71E-37 | 6.24E-37 | 1.493 |
| FX.APH | RLFS | 18473 | 2323 | 1923061 | 1 | 1.25E-66 | 2.02E-66 | 1.498 |
| NM.NT | RLFS | 2111 | 129 | 1606572 | 7.31E-13 | 1 | 1.36E-12 | 0.552 |
| NM.DMSO | RLFS | 3927 | 659 | 1631798 | 1 | 1.22E-34 | 1.82E-34 | 1.746 |
| NM.APH | RLFS | 7002 | 554 | 1996033 | 0.045261 | 0.95877 | 0.089567 | 0.927 |
| FX.NT | DRIP-seq | 4149 | 526 | 642328 | 7.62E-14 | 1 | 1.54E-13 | 0.717 |
| FX.DMSO | DRIP-seq | 9167 | 2658 | 608194 | 1 | 1.07E-155 | 1.55E-155 | 1.914 |
| FX.APH | DRIP-seq | 18473 | 4137 | 634175 | 1 | 1.36E-80 | 2.21E-80 | 1.422 |
| NM.NT | DRIP-seq | 2111 | 389 | 595489 | 0.64755 | 0.37359 | 0.73422 | 1.02 |
| NM.DMSO | DRIP-seq | 3927 | 1229 | 598921 | 1 | 2.21E-90 | 2.98E-90 | 2.083 |
| NM.APH | DRIP-seq | 7002 | 1326 | 641914 | 1 | 1.57E-06 | 2.93E-06 | 1.157 |

P values were calculated using bedtools Fisher's Exact Test. "NT", untreated.

**Table S4.** Number of genes containing DSBs in FX cells that overlap RLFSs or DRIP-seq signals.

| DSB Group | Total DSBs | Genes containing DSBs* | Genes containing DSBs that overlap RLFSs* | Genes containing DSBs that overlap DRIP-seq signals* |
|---|---|---|---|---|
| FXdmso.FXaph.overlap | 6177 (28.8%) | 4121 (27.8%) | 894 (33.1%) | 2364 (37.8%) |
| FXdmso.FXaph.uniquetoFXaph | 12296 (57.3%) | 8715 (58.8%) | 1427 (52.9%) | 3113 (49.8%) |
| FXdmso.FXaph.uniquetoFXdmso | 2984 (13.9%) | 1990 (13.4%) | 378 (14.0%) | 778 (12.4%) |
| FXdmso.FXaph.total | 21457 (100%) | 14826 (100%) | 2699 (100%) | 6255 (100%) |
| NMdmso.NMaph.overlap | 3209 (41.6%) | 2484 (41.4%) | 523 (50.7%) | 1240 (41.3%) |
| NMdmso.NMaph.uniquetoNMaph | 3792 (49.2%) | 2524 (42.1%) | 153 (14.8%) | 718 (23.9%) |
| NMdmso.NMaph.uniquetoNMdmso | 714 ( 9.3%) | 990 (16.5%) | 356 (34.5%) | 1042 (34.7%) |
| NMdmso.NMaph.total | 7715 (100%) | 5998 (100%) | 1032 (100%) | 3000 (100%) |

* Percentages in parentheses indicate the percentage of DSBs or genes in each of the three categories (e.g., XX.XX.overlap, XX.XX.uniquetoXXaph, and XX.XX.uniquetoXXdmso) over the sum of the three categories.

**Table S5. (separate file)**
487 genes that contain co-localized FMRP binding sites and RLFSs (within 1 kb of each other). The table headers are largely self-evident. Columns A through I follow the MACS peak identification output file format (https://github.com/taoliu/MACS#call-peaks). The 487 unique genes are derived from column N ("symbol_g") for those entries for which the value in column AH ("RLFS_1kupdown") is greater than 0.

File attachment: Supplemental_Table_S5.txt

**Table S6.** Select top Gene Ontology terms (p < 0.001) for "Biological pathways" associated with DSBs in the indicated groups.

| DSB Group | Biological pathway | P value | # of genes |
|---|---|---|---|
| FX_UNTREATED | neuron projection development | 1.12E-05 | 100 |
| | nervous system development | 2.08E-05 | 214 |
| | regulation of cell morphogenesis | 2.08E-05 | 66 |
| | synapse organization | 5.29E-05 | 40 |
| | regulation of neuron projection development | 6.25E-05 | 58 |
| | neuron development | 8.09E-05 | 109 |
| | neuron cell-cell adhesion | 0.00015 | 9 |
| | neuron projection morphogenesis | 0.00018 | 70 |
| | ion transmembrane transport | 0.00025 | 107 |
| | inorganic ion transmembrane transport | 0.00048 | 85 |
| FX_DMSO | cell projection organization | 0.00030 | 187 |
| | localization | 0.00030 | 717 |
| | cell part morphogenesis | 0.00050 | 131 |
| | inorganic ion transmembrane transport | 0.00052 | 118 |
| | neuron development | 0.00075 | 146 |
| | cell projection morphogenesis | 0.00086 | 126 |
| FX_APH | positive regulation of GTPase activity | 6.76E-06 | 180 |
| | regulation of cell morphogenesis | 7.35E-06 | 139 |
| | nervous system development | 1.15E-05 | 504 |
| | regulation of cell projection organization | 4.09E-05 | 152 |
| | neuron projection development | 5.56E-05 | 213 |
| | regulation of neuron projection development | 9.09E-05 | 119 |
| | cell adhesion | 0.000107 | 405 |
| | chemical synaptic transmission | 0.000107 | 163 |
| | movement of cell or subcellular component | 0.000199 | 415 |
| | cell projection assembly | 0.000212 | 113 |
| | vesicle-mediated transport | 0.000347 | 343 |
| | synaptic vesicle localization | 0.000569 | 46 |
| | microtubule-based process | 0.000596 | 160 |
| | synaptic vesicle cycle | 0.000681 | 41 |
| | dendrite development | 0.000733 | 63 |
| | cytoskeleton organization | 0.000784 | 271 |
| NM_UNTREATED | None | | |
| NM_DMSO | None | | |
| NM_APH | regulation of cell projection organization | 2.20E-06 | 108 |
| | regulation of cell morphogenesis | 1.56E-05 | 94 |
| | cell projection organization | 3.97E-05 | 201 |
| | regulation of neuron projection development | 0.000217 | 80 |
| | neuron projection development | 0.000559 | 135 |
| | neuron development | 0.000559 | 154 |
| | nervous system development | 0.000646 | 305 |
| | cellular component morphogenesis | 0.000646 | 186 |
| FX_DMSO_RLFS_overlap | flavonoid glucuronidation | 0.000136 | 8 |
| | cellular glucuronidation | 0.000208 | 8 |
| | uronic acid metabolic process | 0.000499 | 8 |
| | flavonoid metabolic process | 0.000499 | 8 |
| | flavonoid biosynthetic process | 0.000499 | 7 |
| | glucuronate metabolic process | 0.000499 | 8 |
| FX_APH_RLFS_overlap | single-organism membrane organization | 0.001285 | 84 |

**Table S7.** Primers for ChIP-qPCR.

| Gene name | Primer Name | Sequence |
|---|---|---|
| ANK1 | ANK1-F | 5'- CTTAGAGGGTGAGGCAGACG-3' |
| | ANK1-R | 5'- ACATCCAGGTGTTTGGCTTC-3' |
| ASTN2 | ASTN2-F | 5'-GTGCTGAGCTTCACACGGTA-3' |
| | ASTN2-R | 5'-AGAGCTGCAGGGTGAACAAT-3' |
| CLNK1 | CLNK-F | 5'-TGTCCCATCTCCTCAGGAAC-3' |
| | CLNK-R | 5'-GCCCAATTCTGCCTCTTTCT-3' |
| DLG1 | DLG1-F | 5'-AGCTTTTCCTTGGAGTGGGTA-3' |
| | DLG1-R | 5'-ATACTTGTGCGGGGGAAGAG-3' |
| GAPDH | GAPDH-F | 5'-GACCTGACCTGCCGTCTAGA-3' |
| | GAPDH-R | 5'-ACCTGGTGCTCAGTGTAGCC-3' |
| GRIA1 | GRIA1-F | 5'-GACCTGACCTGCCGTCTAGA-3' |
| | GRIA1-R | 5'-ACCTGGTGCTCAGTGTAGCC-3' |
| GRM5 | GRM5-F | 5'-GACCTGACCTGCCGTCTAGA-3' |
| | GRM5-R | 5'-ACCTGGTGCTCAGTGTAGCC-3' |
| MTOR | MTOR-F | 5'-GACCTGACCTGCCGTCTAGA-3' |
| | MTOR-R | 5'-ACCTGGTGCTCAGTGTAGCC-3' |
| PTEN | PTEN-F | 5'-GACCTGACCTGCCGTCTAGA-3' |
| | PTEN-R | 5'-ACCTGGTGCTCAGTGTAGCC-3' |
| UGT1A | UGT1A-F | 5'-GACCTGACCTGCCGTCTAGA-3' |
| | UGT1A-R | 5'-ACCTGGTGCTCAGTGTAGCC-3' |
| NC (Chr16:63,868,961-63,869,080) | NC-F | 5'-GACCTGACCTGCCGTCTAGA-3' |
| | NC-R | 5'-ACCTGGTGCTCAGTGTAGCC-3' |

**Movie S1. (separate file)**

Movie for a cell showing co-localization of FMRP and RNA:DNA hybrid.

File attachment: Supplemental_Movie_S1.AVI

#### Supplemental Figure S1

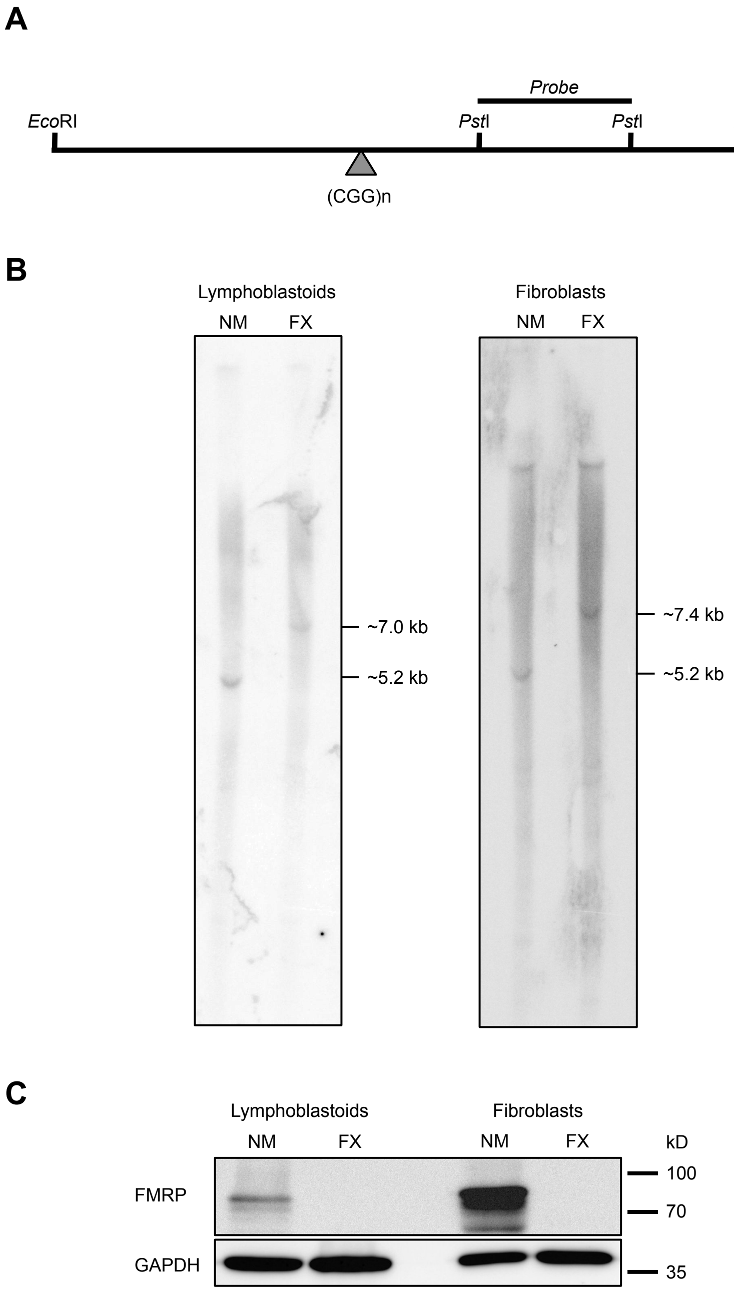

### Supplemental Figure S2

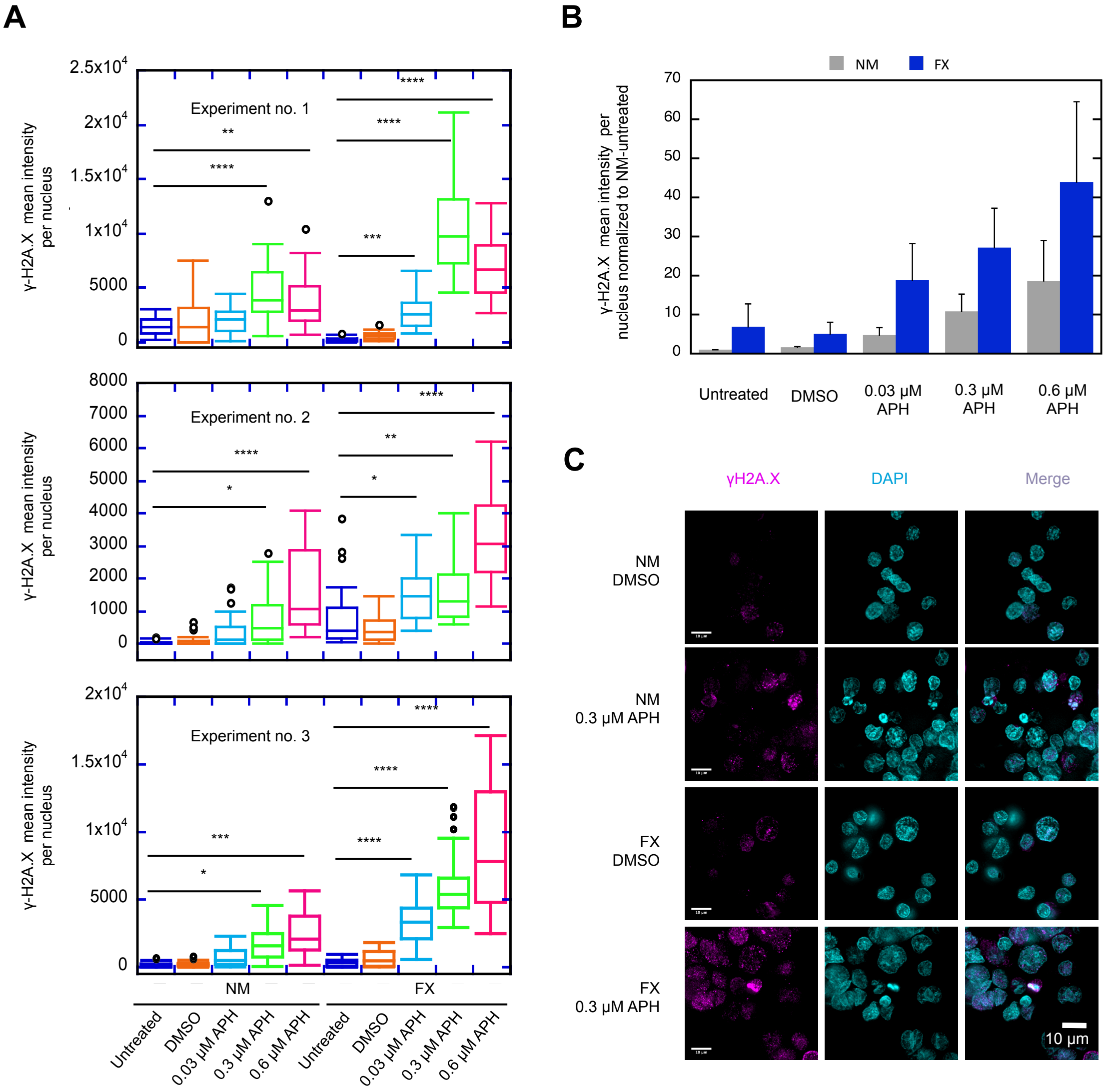

### Supplemental Figure S3

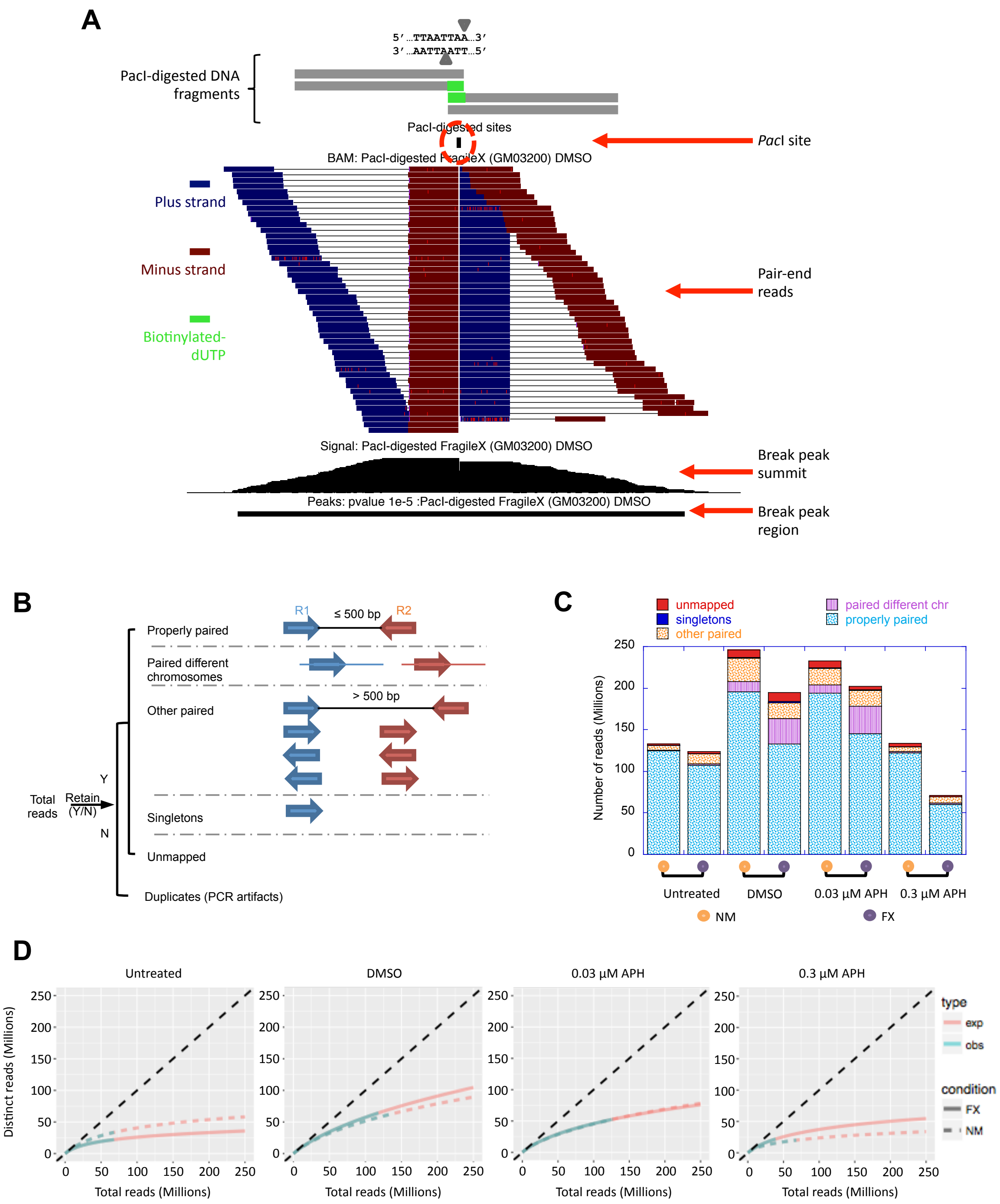

#### Supplemental Figure S4

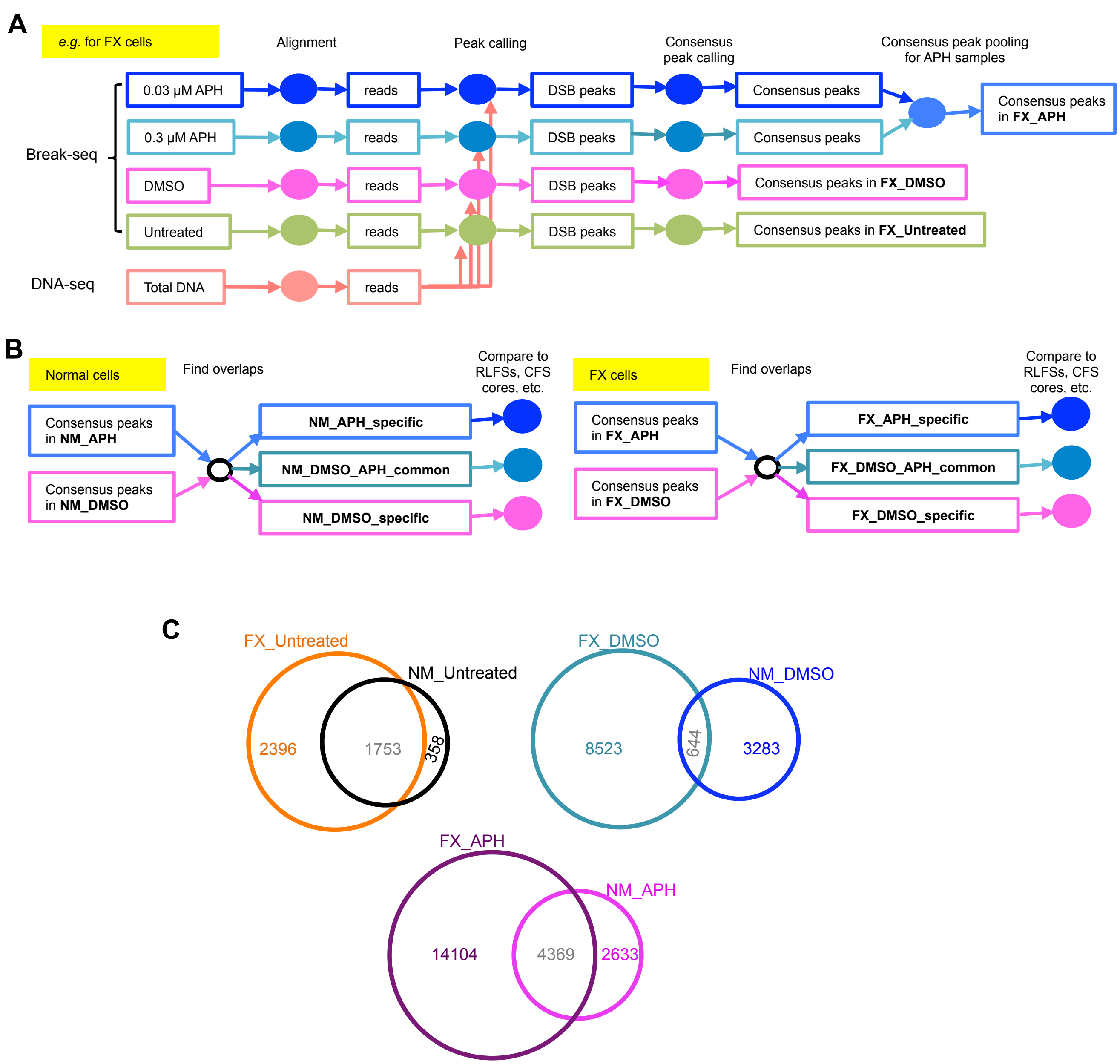

#### Supplemental Figure S5

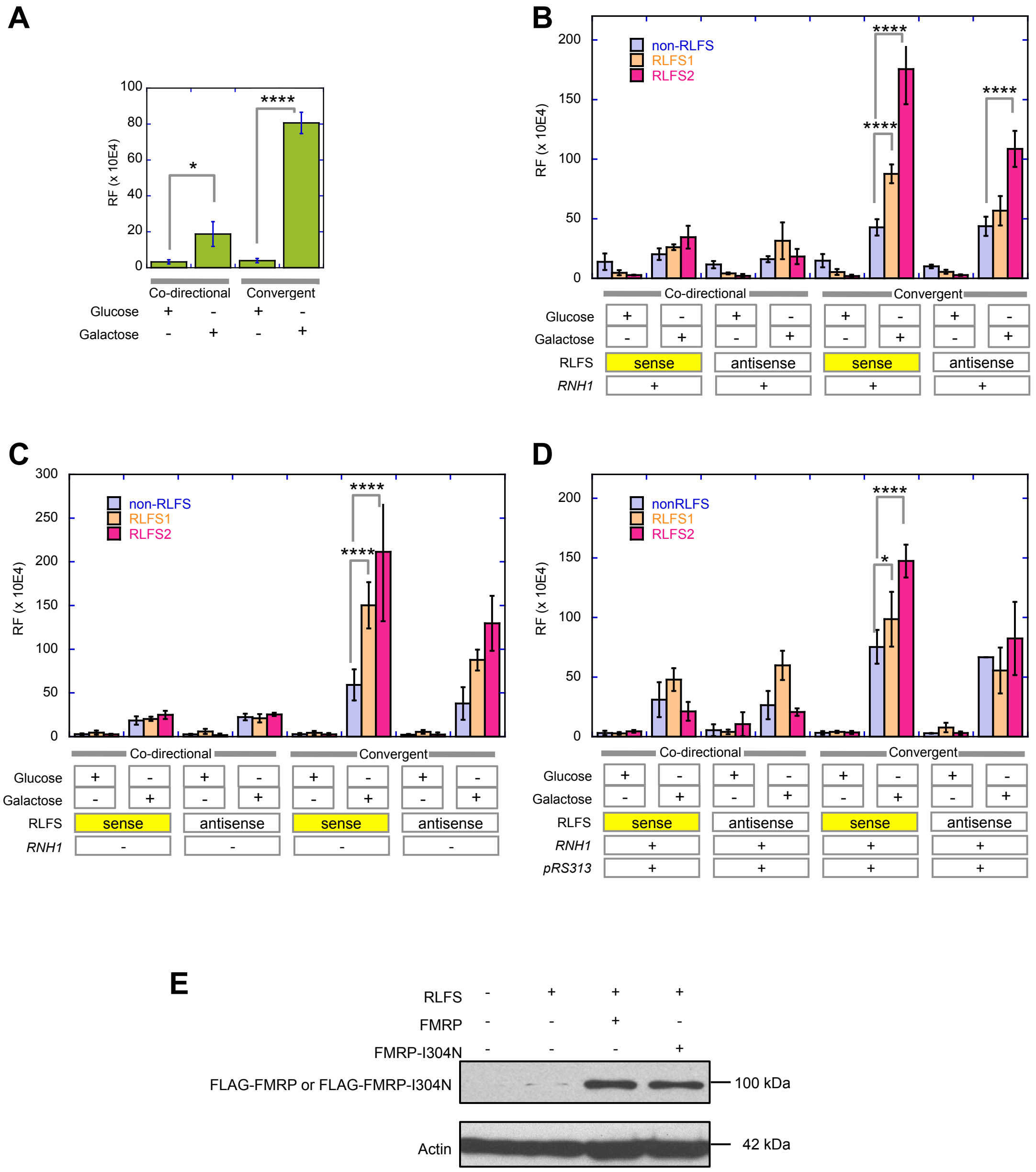

#### Supplemental Figure S6

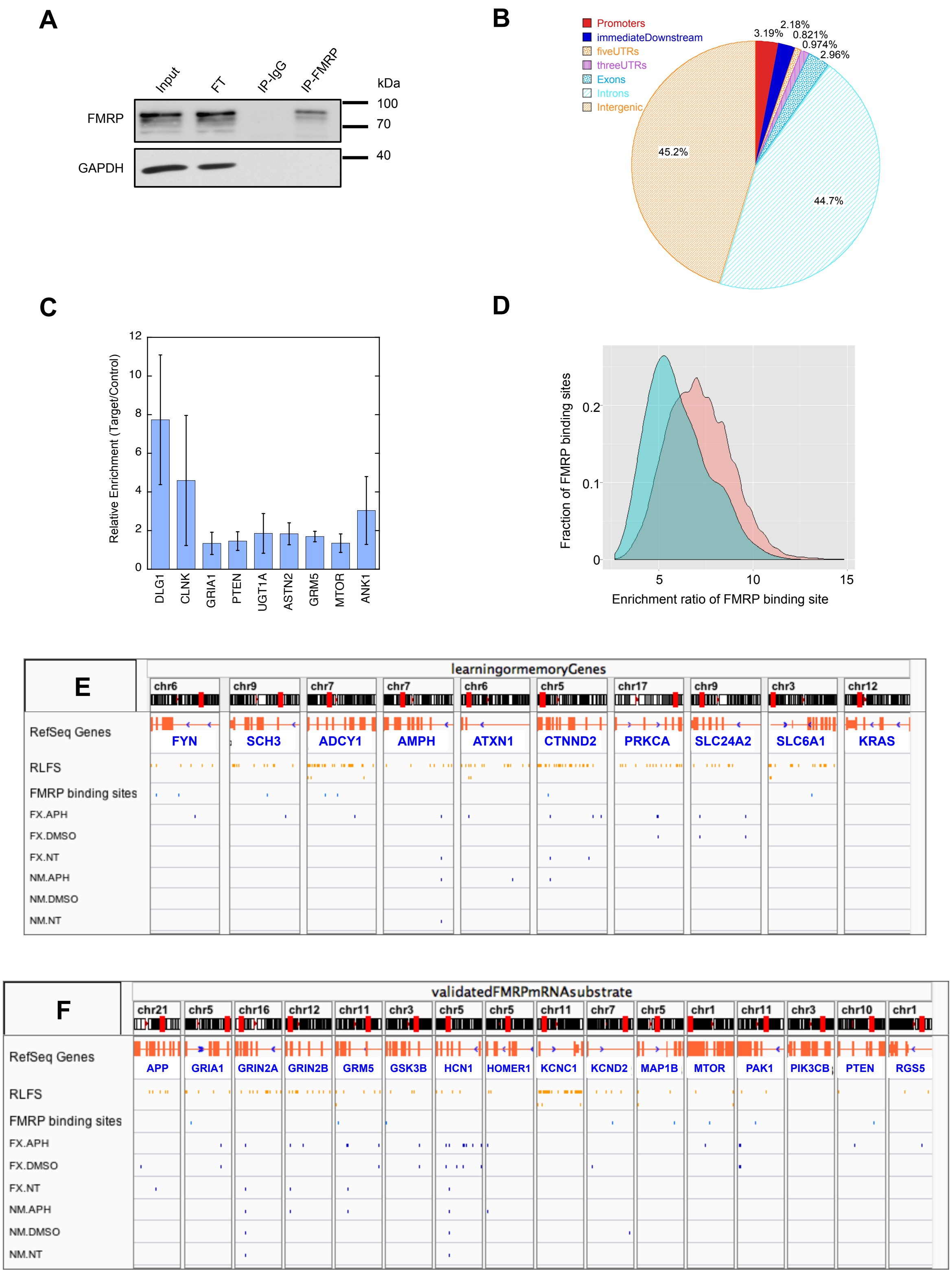

#### Supplemental Figure S7

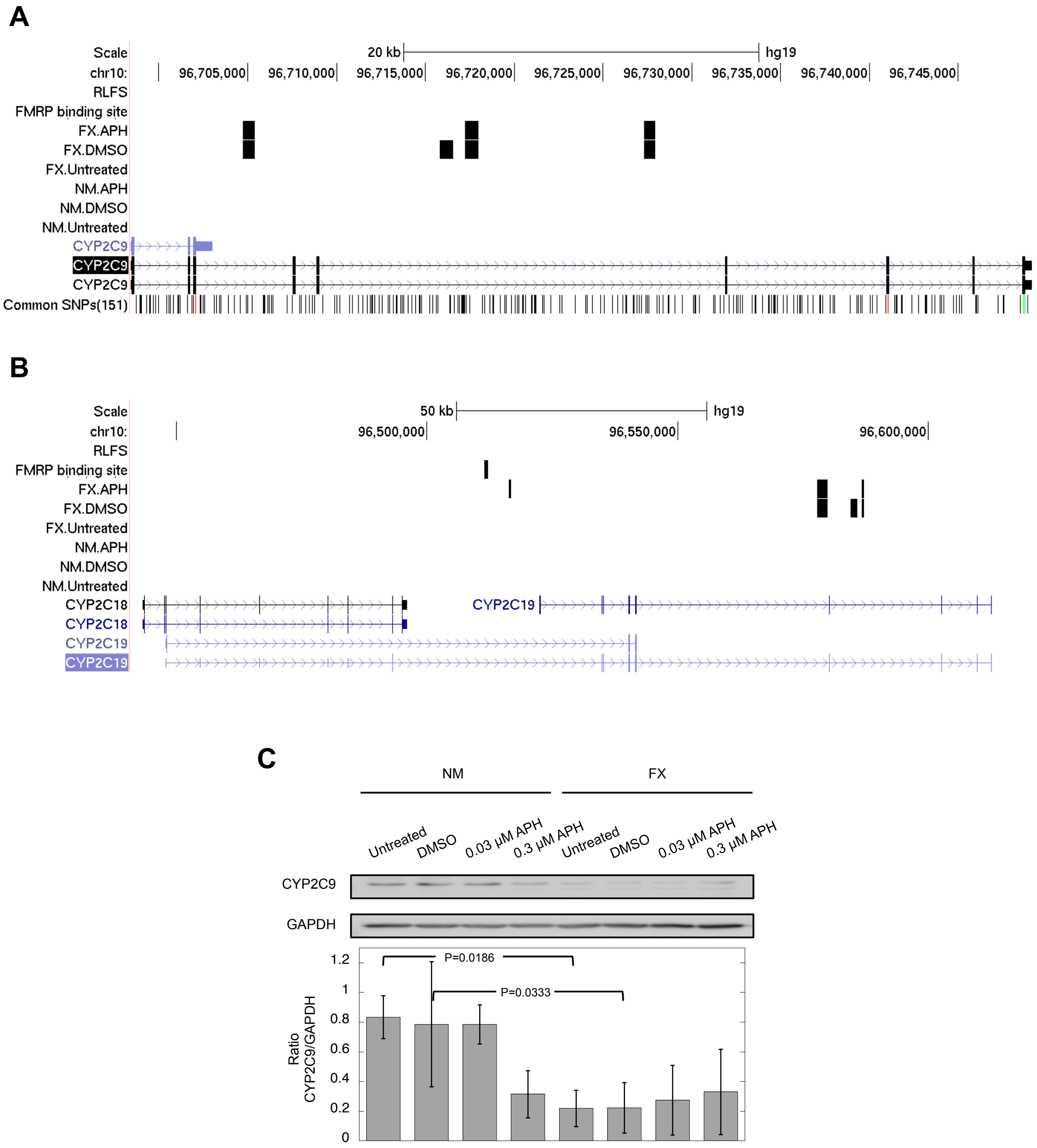

### Supplemental Figure S8

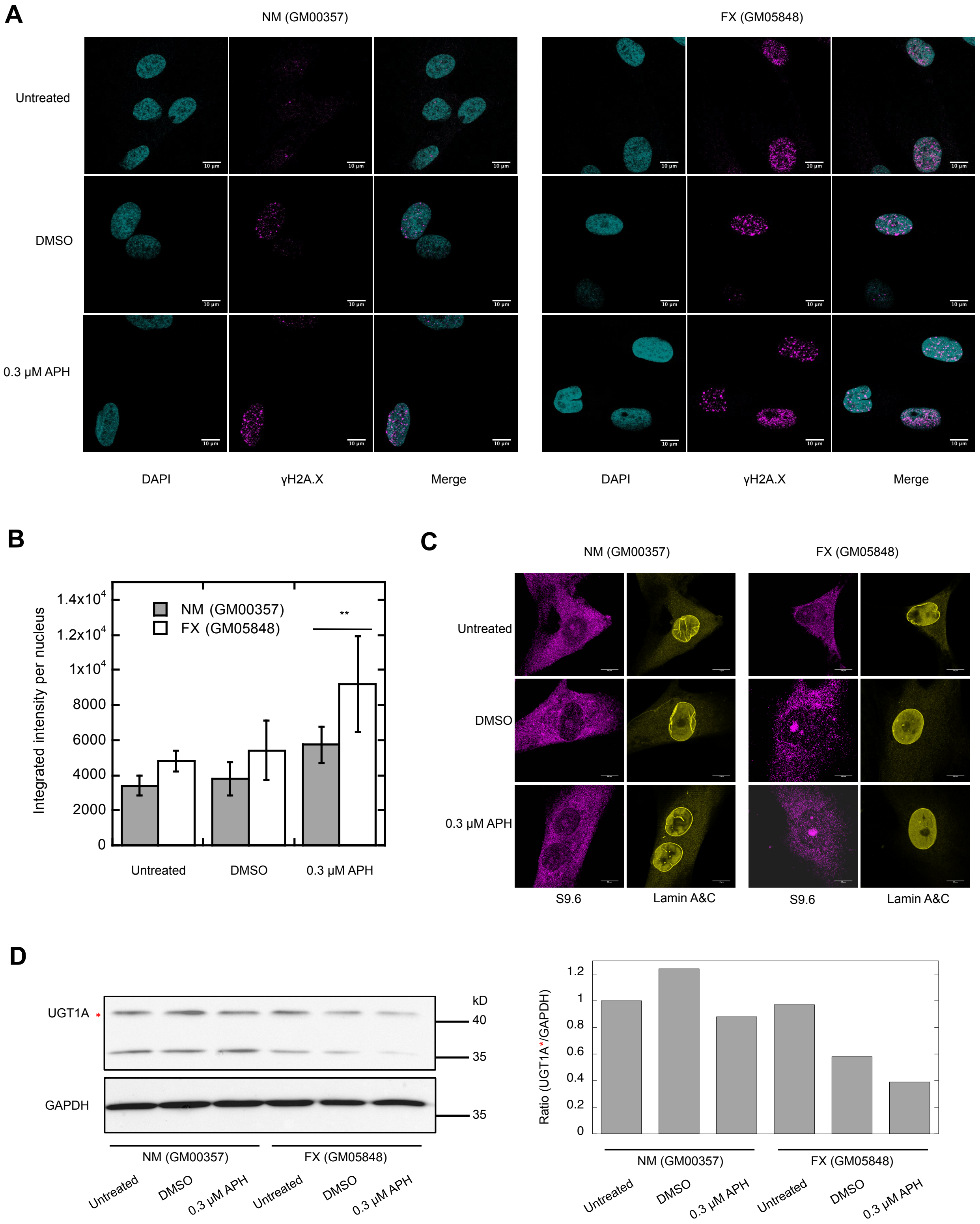

### Supplemental Figure S9

**A**

| Sample | Whole genome (Obs/Exp) | Early Replicating (Obs/Exp) | Late Replicating (Obs/Exp) |
|---|---|---|---|
| NM.Untreated | 141/106 | 33/26 | 100/85 |
| NM_DMSO | 337/195 | 44/56 | 265/143 |
| NM_APH | 568/333 | 75/79 | 456/273 |
| FX_Untreated | 226/200 | 49/51 | 171/162 |
| FX_DMSO | 781/427 | 155/106 | 566/332 |
| FX_APH | 1219/849 | 283/221 | 854/635 |

**B**

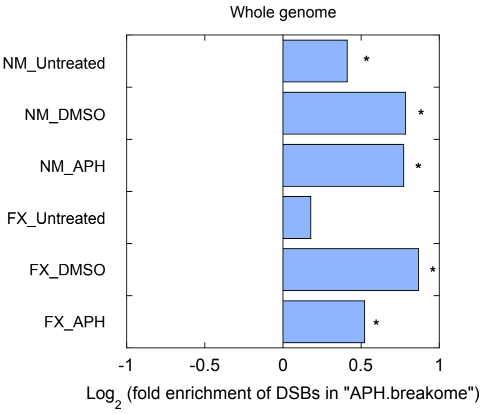

**C**

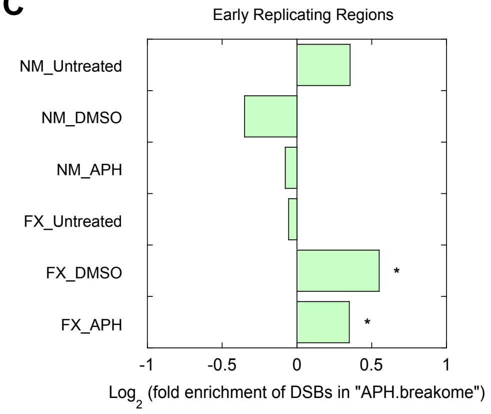

**D**

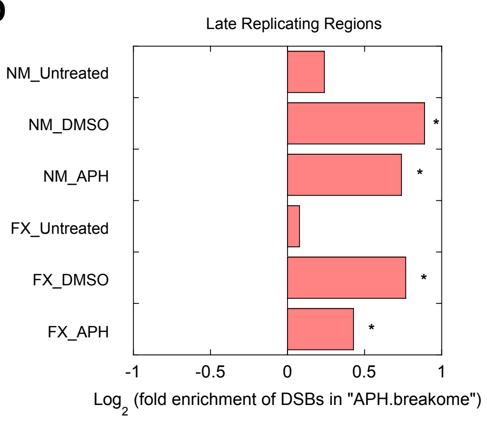
